# Design and Assembly of a Cargo-agnostic Hollow Two-lidded DNA Origami Box for Drug Delivery

**DOI:** 10.1101/2024.03.27.586853

**Authors:** Abigail Koep, Nabila Masud, Jaylee Van’t Hul, Carson Stanley, Marit Nilsen-Hamilton, Anwesha Sarkar, Ian C. Schneider

## Abstract

DNA origami, a method of folding DNA into precise nanostructures, has emerged as a powerful tool to design complex nanoscale shapes with movable parts. DNA origami has great potential as a drug delivery system that can encapsulate and protect a range of cargos spanning small molecules through large proteins, while remaining stable in a variety of *ex vivo* processing conditions and *in vivo* environments. DNA origami has been utilized for drug delivery applications, but the vast majority of these structures have been flexible, flat 2D or solid 3D nanostructures. There is a crucial need for a hollow and completely enclosed design capable of holding any type of cargo. In this paper, we present the design and assembly of a hollow DNA origami “box” with two actuatable lids. We characterize isothermal conditions for structural assembly in minutes that eliminates the need for a thermocycler. The stability of these structures is outstanding, remaining stable at body temperature and low pH for weeks and in the presence of solvents and biological fluids over several days. We demonstrate that passive loading of small molecules is charge dependent. We also outline an approach to design staple extensions pointing into the cavity or outside of the hollow DNA origami, allowing for either active loading of protein or the potential for decoration with passivating or targeting molecules. Future work includes fitting this hollow DNA origami structure with alternative lid opening mechanisms to release a variety of different cargos in response to environmental cues.

**Graphical Abstract:** 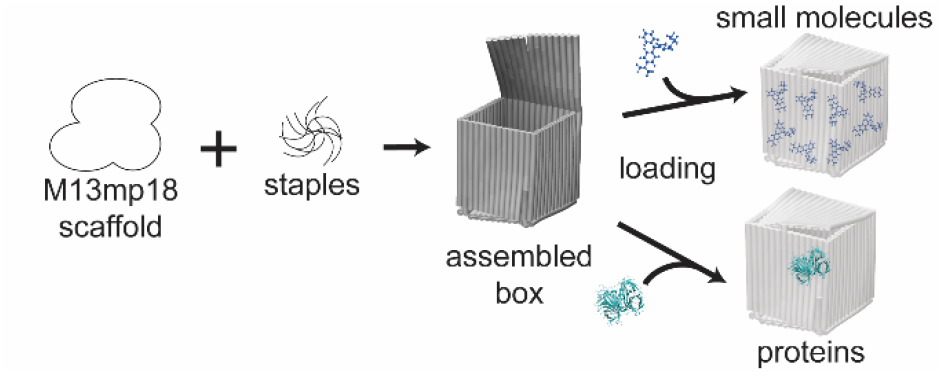

## 1. Introduction

The need for nanoparticle drug delivery systems is growing. Several key features are important to consider when designing these systems. They should be able to encapsulate, retain and protect the drug from degradation or improperly timed release. They should be able to target particular parts of the body and release drugs gradually over time. Ideally, this controlled release would be in response to environmental cues, potentially even performing logic operations on multiple environmental signals. Other notable features include ease of assembly, stability in biological environments and tunable immunogenicity.^1–3^ Several types of nanoparticles have been developed from diverse materials, such as lipids, polymers, or inorganic substances for various uses.^4^ While useful in numerous applications these systems lack an ability to finely tune nanoparticle structure. DNA origami presents a nanoparticle system with a tremendous ability to finely tune shape as well as generate mechanically sensitive or movable components. DNA nanostructures are being utilized across several fields from drug delivery to sensors to mechanical devices.^5–14^

DNA origami is a method of folding DNA into specific shapes.^15^ The single-stranded DNA (ssDNA) sequence called the “scaffold” provides a backbone for the designs, while shorter ssDNA strands called “staples” hold the design in place. DNA origami has tremendous potential for drug delivery due to its ability to form a variety of mechanically movable shapes, its stability, non-immunogenic properties and ease in chemically functionalizing the structure for various purposes.^3,10,16–18^ For DNA origami drug delivery constructs to be a viable option in medicine, they need to be designed to encapsulate and confine a variety of different cargos that can be released in response to environmental cues. Additionally, the assembly and purification methods need to be optimized for ease of production and separation from excess staple strands. This has proven challenging as most protocols for DNA origami assembly have required changes in temperature over long time periods during assembly, which limits production to small volumes assembled in a thermocycler.^19–21^ There are some examples of isothermal assembly, but incubation times are long.^19,21^ DNA origami have been shown to be reasonably stable, but this seems to depend highly on the shape of the structure,^22,23^ necessitating an examination of the stability for each structure.

The vast majority of DNA origami structures are flexible, flat 2D structures of different geometries. These have been used to deliver or release drugs or model molecules including doxorubicin, platinum-based molecules, and antibody fragments to name a few.^8,10,12,24–26^ Molecules have been attached to the surface of the DNA origami structure through electrostatic interactions as well as through covalent linkages. For instance, doxorubicin and similar molecules have been bound electrostatically to the surface of a 2D DNA surface.^12,13,24^ Folic acid and other molecules have been covalently linked to similar 2D DNA origami shapes.^10^ 2D DNA origami structures are easy to design, assemble, and functionalize compared to 3D structures, but 2D structures have limitations. Due to the nature of “loading” cargo on the structures, they are unable to provide protection or controlled release, because the cargo is not completely enclosed. It also is challenging to encode sensitivity to environmental cues in 2D structures, whereas 3D DNA origami structures can be designed to specifically to respond to environmental signals.^27,28^

3D DNA origami structures have potential to encapsulate cargo completely with release mechanisms for environmental cues. DNA origami box or cube designs with lids that can open in response to logic nucleic acid keys have been fabricated.^27,29^ While these boxes were shown to open on demand, their loading with cargos was not demonstrated. Other DNA origami designs show hollow tubes or wireframe structures that can gradually release drug when delivered, but the release is uncontrolled due to the open cage designs.^2,3,12,25^ Another design is a DNA origami tube lacking lids that can be opened along its length based on logic nucleic acid keys (similar to what was mentioned above). Each of these tube designs still suffer from non-specific delivery due to open-ended or wireframe designs.^2,3,8,25^ Enclosed 3D DNA origami structures have been designed with pH responsive lids.^30^ However, the loading potential is minimal in this design due to edges that are multiple helices thick, resulting in a small internal volume.^30^ There is a need for actuatable “box” designs that can be loaded with sufficient cargo and for which numerous alternative release mechanisms in response to specific environmental cues could control delivery.

Our long-term goal is to design 3D hollow DNA origami structures to entrap and protect cargo that can later be released in response to a variety of specific environmental cues. To address this need, hollow DNA origami boxes were designed with two actuatable lids that allow loading and release of a cargos ranging from small molecules to large proteins. We used Atomic Force Microscopy (AFM) and Dynamic Light Scattering (DLS) to characterize the newly designed DNA origami structures. We developed an isothermal assembly approach that dramatically diminished the assembly time and optimized purification methods for these hollow DNA origami boxes. Stability was characterized under relevant processing and biological conditions, including various temperatures and pH values, as well as in the presence of solvent and serum. We demonstrated loading of various cargos spanning small molecules to proteins. We used both passive loading as well as active loading, leveraging a design approach for determining optimal staple extension length and position that allows for protein entrapment within the DNA origami box or the decoration of protein on the outside decoration of the DNA origami box. In this article, we outline the design, fabrication, and characterization of a hollow DNA origami box for drug delivery with two actuatable lids for potential release by two distinct environmental cues.

## 2. Results and Discussion

### 2.1 Design, Assembly, and Purification

#### 2.1.1 Design and Temperature Gradient Assembly

A DNA origami design with multiple lids can allow for multiple responses to environmental cues and enhance molecular transport into or out of the hollow DNA origami structure. Cadnano (square) was used to design the DNA origami box with two actuatable lids (**Supplemental Figure 1**). Staple sets are available in supplementary information (**Supplemental Figure 2 and Supplemental Table 1**). The DNA origami box consists of an uninterrupted squared-shaped conduit with two hinged lids that can attach to the respective opposite edges of the conduit to close the box. DNA helices can be flexible, so to create stable edges in the design, the DNA staples crossed helices at the edge. This ensures all the helices are connected at the edge.^31^ The design is for a box that is 40x40x40 nm in size, with a diagonal of approximately 56 nm. Cadnano designs with the closed DNA origami box, as well as versions with one lid or two lids open were rendered using CanDo and UCSF ChimeraX (**Figure 1A**). The size for each design was measured using Dynamic Light Scattering (DLS) (**Figure 1B**). Closed boxes were smaller than either one-lid-open or two-lids-open versions, matching our expectations due to the flexible nature of the lids. To visualize individual structures, Atomic Force Microscopy (AFM) images were taken of each design (**Figure 1C**). These hollow DNA origami boxes are fragile and collapse and aggregate when dried unlike most 2D DNA origami structures, so in-solution AFM was used to visualize the boxes (**Figure 1C**). The estimated sizes matched our expected size based on both Cadnano predictions and DLS measurements. The DNA origami boxes with closed lids, one-lid-open, or two-lids-open showed characteristically distinct shapes (**Figure 1D**). Closed boxes were symmetric in shape. One-lid-open boxes were frequently asymmetric with one side with a shorter height (the lid) than the other (the box).

**Figure 1:**
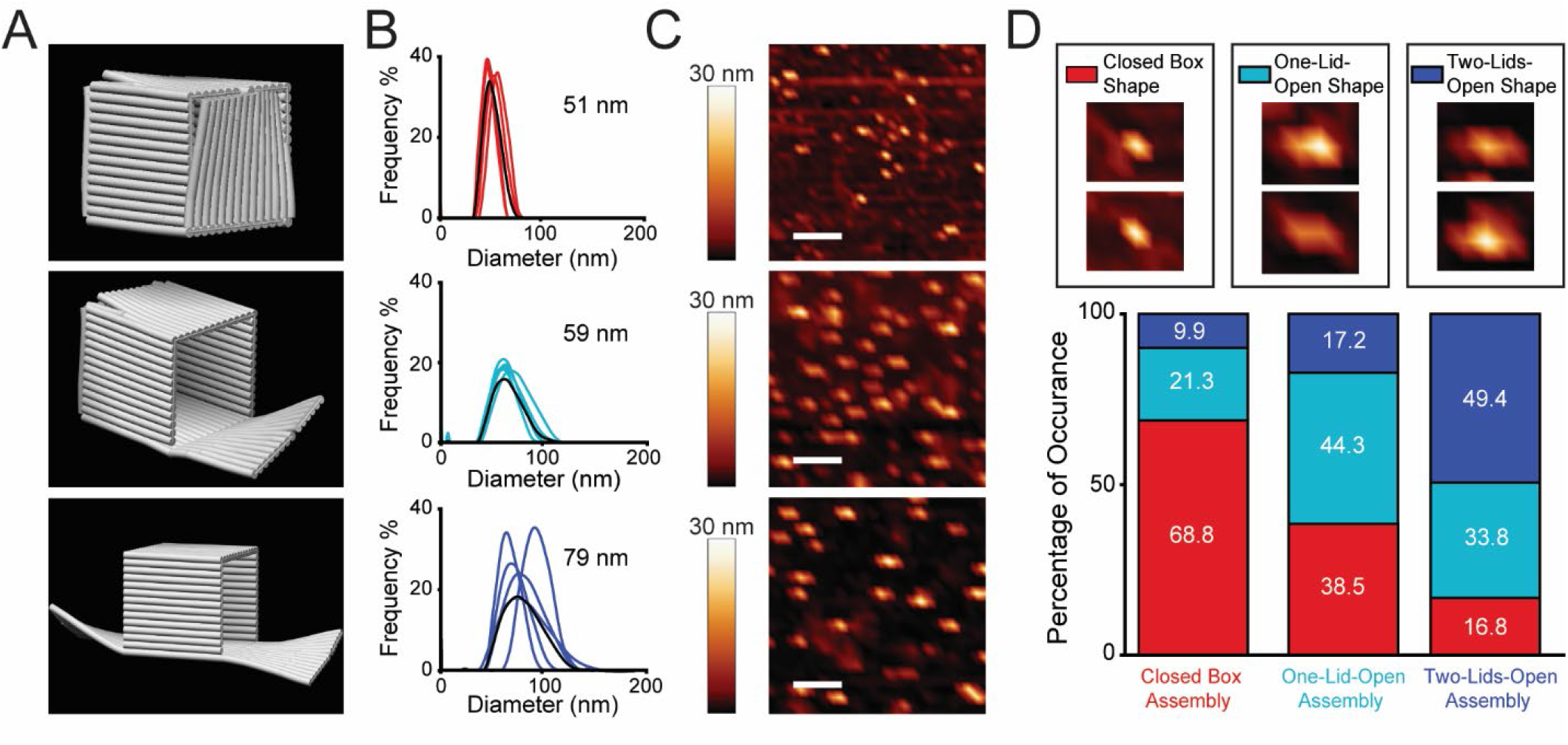
Characterization of hollow DNA origami boxes with two openable lids. A) CanDo renderings of cadnano files based on the design of the DNA origami box with no lids open, one-lid-open, and two-lids-open. B) DLS graphs of the size distribution after gradient-temperature assembly and filter purification. The average diameter distribution from five independent assemblies is noted as the black curve and the peak diameter is denoted on the graph. C) AFM images of each gradient-temperature assembly (purified at 5k x *g* for 20 min, 3 μM DNA origami). Scale bar represents 200 nm. D) Shape quantification of the AFM images with sample images to for each expected assembly shape. Counting was performed on four AFM images from each of two independent assembly experiments per design. The graph shows the fraction of the shapes within an assembly.

Two-lids-open boxes were symmetric or asymmetric, but usually contained a tall structure in the center (the box) with two shorter structures (the lids) on the sides. Quantification of the shapes representing each structure in each assembly (closed box, one-lid-open and two-lids-open) showed that the largest fraction of structures for each assembly was the predicted shape (**Figure 1D**). These results showed that we can assemble hollow DNA origami boxes that are closed or that have either one lid or two lids open using traditional temperature gradient assembly.

#### 2.1.2 Isothermal Assembly

Given we were able to assemble DNA origami boxes using a typical gradient assembly, we were interested in whether we could use other assembly methods to fabricate hollow DNA origami boxes. As demonstrated, hollow box DNA origami assembly is relatively easy in small quantities needed for structural characterization, but larger quantities needed for *in vivo* experiments or even *in vitro* drug release experiments are difficult to make because of the complex assembly requirements. DNA origami is nearly always assembled with a thermal gradient of -1°C/min from 95 °C to room temperature that is accomplished in a thermocycler in 50 µL aliquots. This tedious process wastes time and material.

Therefore, we explored whether isothermal assembly, which allows for larger batch production, could produce these hollow DNA structures. Hollow DNA origami structures were assembled for 2 h at a range of assembly temperatures from 4 °C to 90 °C after heating the assembly mixture to 95 °C for 10 min, the samples were resolved by electrophoresis through an agarose gel (**Figure 2A**). Normal gradient assembly results in a band that runs slower than the scaffold (∼5000 bp). In addition, there is a weak secondary band that runs more slowly (10,000 bp to 20,000 bp). High temperatures (> 80°C) resulted in a smear in the lane instead of a clear band for the assembled DNA origami boxes, whereas lower temperatures produced a band at the same size as that produced using the temperature gradient assembly method. Isothermal assembly resulted in DNA origami structures with similar sizes to those achieved with a thermal gradient as measured by DLS and AFM (**Supplementary Figure 3**). These results suggested that hollow DNA origami structures are produced during isothermal assembly (**Figure 2A**).

**Figure 2:**
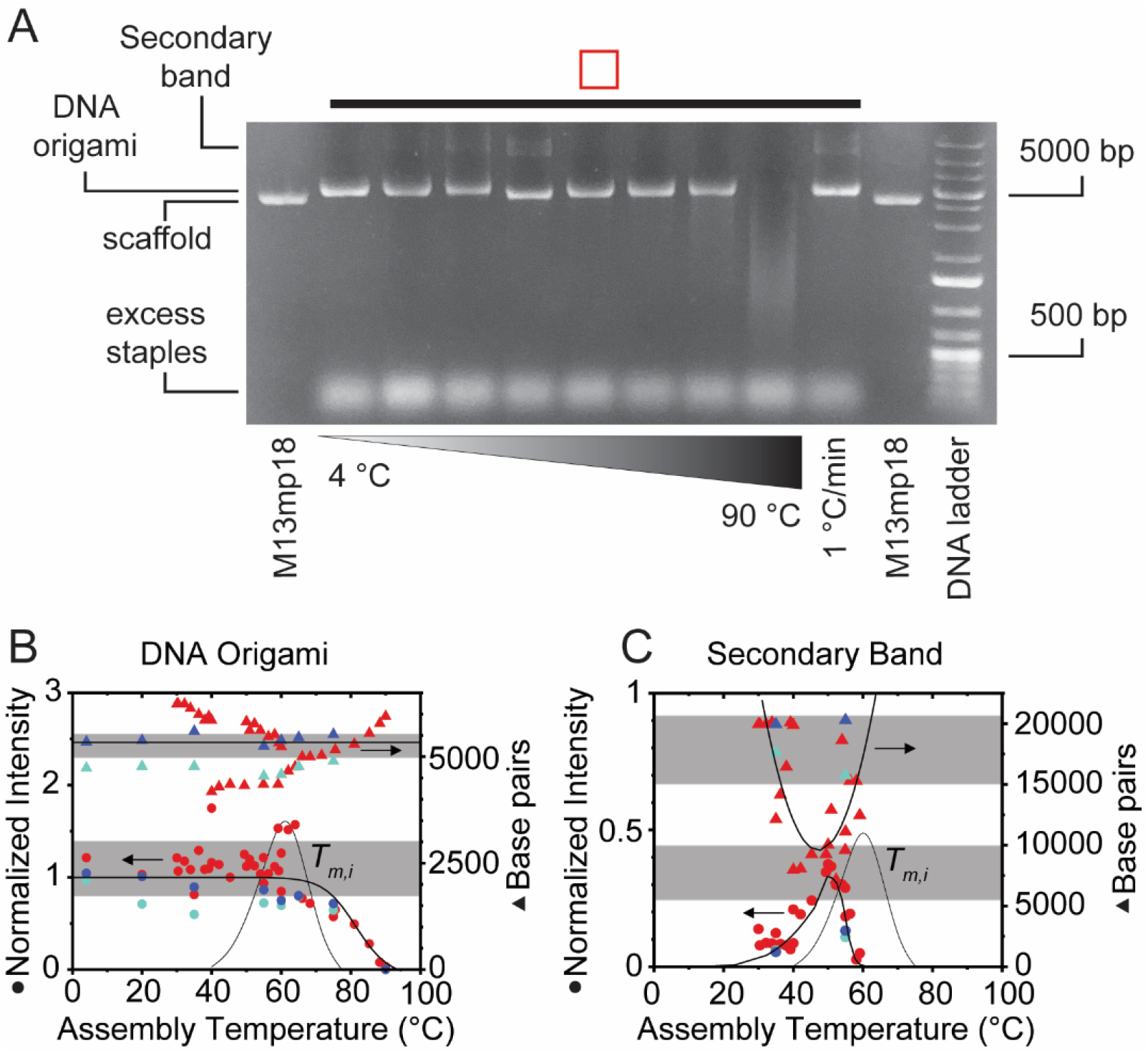
Temperature-dependent assembly of hollow DNA origami boxes under isothermal conditions. A) Agarose gel of different assemblies over a range of temperatures after heating the assembly mixture for 10 min at 95 °C. Band intensity and apparent size of the DNA origami assembly B) and the DNA band that runs more slowly than the DNA origami assembly C). B) The triangles represent apparent size in terms of base pair number as measured relative to the DNA ladder. The circles represent normalized fluorescent intensity of the bands. Red is the closed box assembly. Light blue is one-lid-open assembly. Dark blue is two-lids-open box assembly. The grey shaded regions are the observed intensity/size ranges when assembly is done using a thermal gradient of -1 °C/min starting from 95°C. Solid lines are melting temperature isotherm^48^ of complementary DNA binding fit to the data. The thin line shows the distribution of melting temperatures of the staple strands.

To better characterize the isothermal assembly, we quantified both the band intensity and apparent DNA size across numerous isothermal assembly temperatures and plotted the distribution of melting temperatures of staple strands (*T_m_*_,*i*_) on the same graph (**Figure 2B**). Isothermal assembly temperatures below the majority of the individual staple strand melting temperatures result in hollow DNA origami assembles with the same intensity and size that we see with the gradient assembly (**Figure 2B**). No difference was seen between closed box, one-lid-open boxes or two-lids-open boxes in their isothermal assembly. Others have shown isothermal assembly for simple 2D and non-hollow 3D DNA origami designs using a handful of annealing temperatures from 15-60 °C with annealing times ranging from 5 min at lower temperatures to 4 h at higher temperatures.^19,21^ Modeling has shown a thermodynamic preference for DNA origami annealing at temperatures less than the staple strand melting temperatures.^32^ In addition to the primary band, we observed a secondary band. A similar temperature dependence indicative of isothermal assembly was observed with the secondary band normalized intensity and apparent size. Normalized intensity peaked at around 50 °C, while apparent size was a minimum at a similar temperature (**Figure 2C**). The maximum and minimum were at a slightly lower temperature than the peak in staple strand melting temperatures. This secondary band could be dimerized boxes or structures with incomplete assembly. At high temperatures, the dimerization could be inhibited due to higher internal energy, whereas at lower temperatures the number of misfolded structures due to poor base pairing could be inhibited. We assessed the kinetics of isothermal assembly with an eye toward diminishing the incubation time.^19,32^ Amazingly, hollow DNA origami structures assembled very quickly on the order of a minute or so between the temperatures of 4 °C and 90 °C (**Supplemental Figure 4**).

DLS confirmed that the DNA origami boxes are present in solution after only 10 min of assembly at 4 °C (**Supplemental Figure 3A,B**).^33^ These results show that we are able to assemble hollow DNA origami structures isothermally on the order of minutes if the assembly temperature is below a large fraction of the staple strand melting temperatures.

#### 2.1.3 Purification

When assembling DNA origami, excess staples are used to ensure complete assembly. After assembly, purification of DNA origami by removal of excess staples eliminates the chance of staples interfering with characterization assays. In these studies, purification was done using concentrating filters with a MWCO of 100k. Based on the filter’s recommendations, the hollow DNA origami was filter purified from staples using 20k x *g* centrifugal force for 5 min. However, this did not remove the excess staples from preparations of closed boxes, one-lid-open boxes, or two-lids-open boxes (**Figure 3A**).

**Figure 3:**
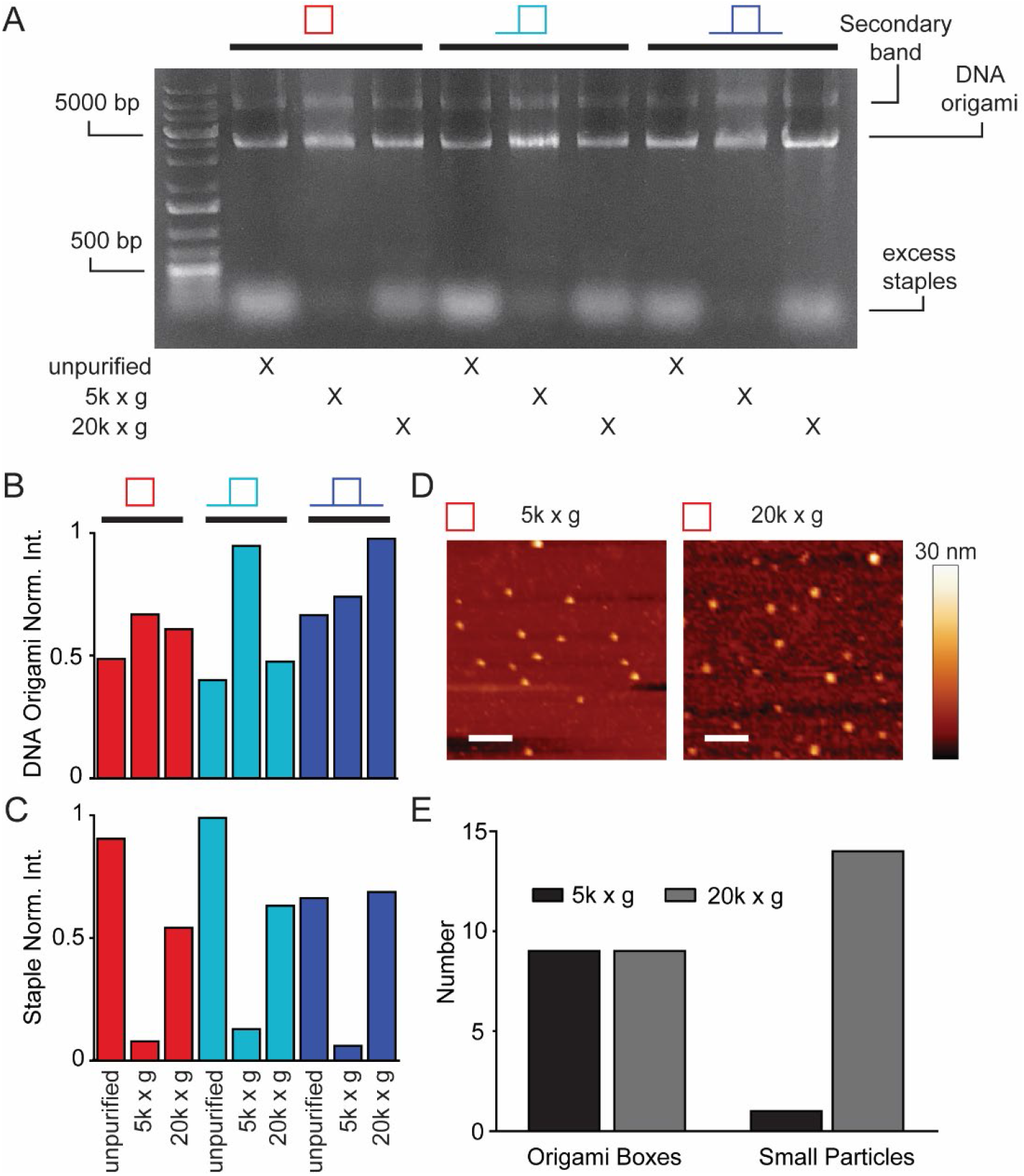
Hollow DNA origami box separation is sensitive to centrifugal force. A) Agarose gel showing the lack of staples when the assemblies are purified at 5k x *g* for 20 min compared to no purification at 20k x *g* for 5 min. B) DNA origami and C) normalized staple band fluorescence intensities for closed box assemblies (red), one-lid-open box assemblies (light blue) and two-lids-open box assemblies (dark blue) shown in A). Intensities were normalized to the largest fluorescence intensity. D) AFM images of the origami structures obtained after purification at each of the two centrifugal forces (3 μM DNA origami) and E) quantification of number of DNA origami boxes vs. small box fragments. Counting was performed on one AFM image from a single gradient-temperature assembly experiment per condition. Scale bar represents 200 nm.

Concerned that we were crushing the structures, we decreased the centrifugal force to 5k x *g* for 20 min to minimize its effects on these fragile hollow DNA origami boxes.^34^ At this lower speed and longer run time, the hollow DNA origami structures were retained and excess staples were successfully eliminated (**Figures 3B&C**). To determine if there was damage to the boxes, we imaged them using AFM (**Figure 3D**). We observed similar numbers of hollow DNA origami boxes by AFM but more staples (small particles < 30 nm) after separation by 20k x *g* compared with 5k x *g* (**Figure 3D&E**). Failure to remove excess staples seems to be specific to these hollow DNA origami boxes as 2D DNA origami structures can be purified at high centrifugal force. We propose that the higher spin speeds crush a small fraction of hollow DNA origami on the surface of the filter. This clogs the filter and eliminates the ability to filter the excess staples at high spin speeds. However, lower spin speeds do not crush the boxes and thereby allow for the removal of more excess staples. Consequently, we have identified proper purification conditions that allow for excess staple removal from these hollow DNA origami boxes.

### 2.2 Stability

Once we were able to assemble and purify the hollow DNA origami structures, we were interested in examining their stability. Stability is an important aspect of drug delivery systems. For DNA origami to be used in drug delivery, it should be stable in different environments. Temperature stability will determine if refrigeration is needed for storage and if the drug delivery construct will disassemble at body temperature. Stability at different pHs will determine its potential for application in different tissues or in different compartments within the cell like lysosomes, where acidic conditions are present. Finally, DNA origami drug carriers need to be compatible with other components of the drug formulation, such as solvents like dimethyl sulfoxide (DMSO), and with biomolecules present in body fluids. We have modeled the latter with fetal bovine serum (FBS). Our base case was DNA origami assembled in the refrigerator (4 °C) at neutral pH (7.5) with 12.5 mM Mg(OAc)_2_ in the absence of DMSO and FBS. To test stability, we altered one condition from the base case, probing stability of the DNA origami boxes at different temperatures (4 °C, 18 °C, and 37 °C) and pH values (pH 5 and pH 7.5) for up to 14 days and in solvents (10% DMSO) and biofluids (10% FBS) for up to 3 days. After each incubation time, samples were collected and assessed using electrophoretic resolution through agarose. The base case showed no change in either apparent size or band intensity (**Figure 4A, B, E&F**). Altering the temperature to 18 °C resulted in no change in apparent size over the entire incubation time (**Supplemental Figure 5A**), but a slight increase in intensity, however the confidence intervals indicate that this is likely not significant.

**Figure 4:**
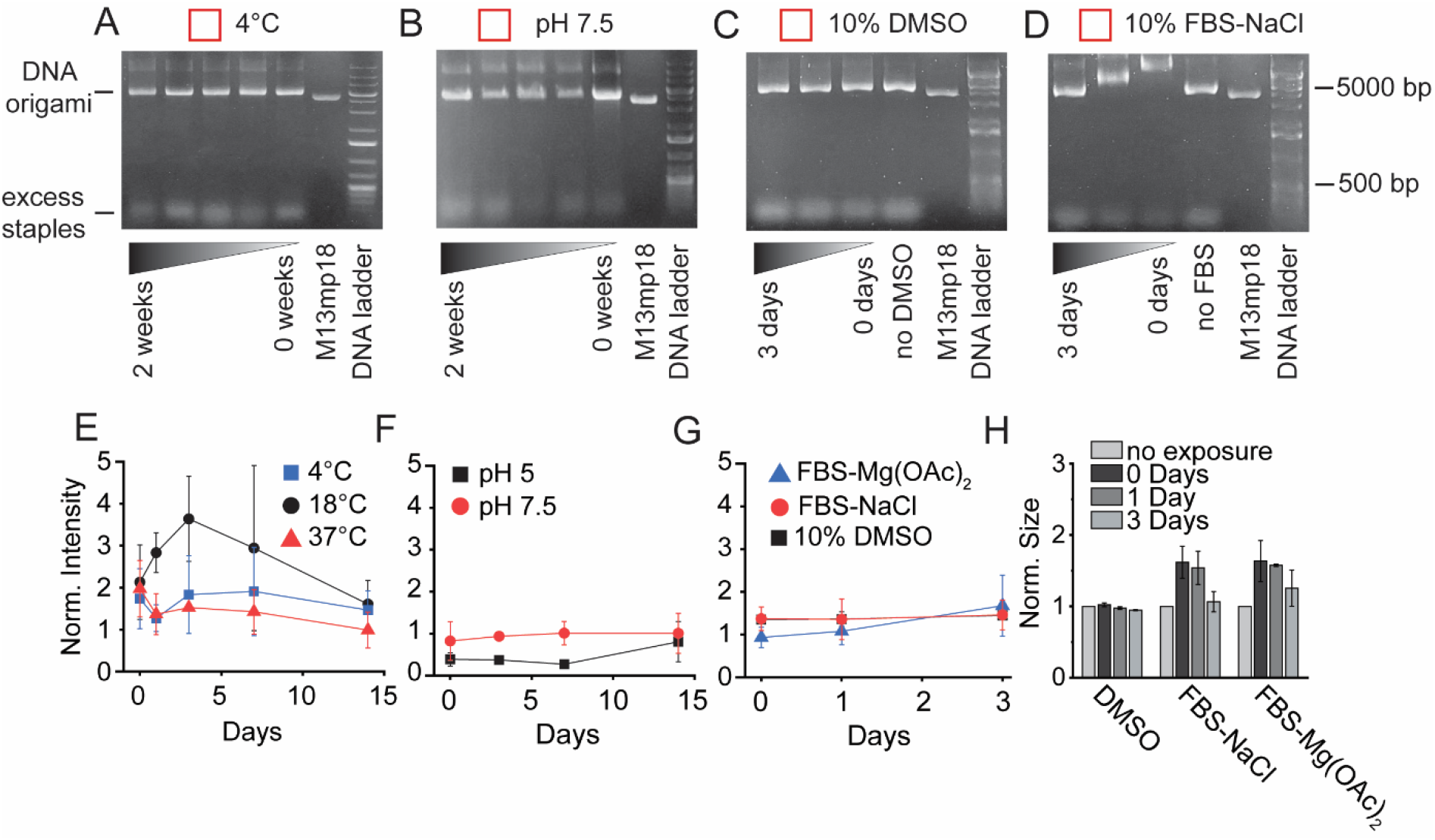
Hollow DNA origami boxes are temperature, pH and solvent stable. Electrophoretic resolution through agarose gels of hollow closed box DNA origami isothermal assemblies held for various times A) at 4 °C, B) at pH 7.5, C) in 10% DMSO and D) assembled in 100 mM NaCl and incubated in 10% FBS. Quantification of the DNA origami band intensity for DNA origami assemblies kept at different E) temperatures, F) pH, G) solvent or serum conditions. Each dot represents the average of three independent assembly experiments. H) Quantification of the DNA origami band size in base pairs normalized to the no serum exposure DNA origami band size as measured in base pairs. DNA origami assemblies kept in 10% DMSO, assembled in 12.5 mM Mg(OAc)_2_ and incubated in 10% FBS and assembled in 100 mM NaCl and incubated in 10% FBS. Each bar represents the average of three independent assembly experiments. All error bars represent 95% confidence intervals.

Higher temperatures (37 °C) showed neither a change in apparent size or intensity over the entire incubation time. Altering the pH to 5 or adding 10% DMSO also did not change the apparent size or band intensity over the entire incubation time (**Figure 4C, F&E** and **Supplemental Figure 5B**). This suggests that these boxes are stable to temperatures as high as body temperature and as low as pH 5 for at least two weeks and in the presence of 10% DMSO for at least three days.

Previous studies have reported that assembly in other salts results in enhanced DNA origami stability.^35,36^ We probed whether these hollow DNA boxes can be assembled in sodium chloride in the absence of magnesium salts. No dramatic changes were observed when the hollow DNA origami boxes were assembled in sodium chloride (**Supplemental Figure 5C** and **Figure 4D**). The most stringent test for stability is in 10% FBS. Serum contains nucleases that can degrade DNA origami boxes.^37,38^ We examined the stability of hollow DNA origami boxes assembled in either Mg(OAc)_2_ or sodium chloride and exposed to 10% FBS (**Figure 4D, G&H** and **Supplemental Figure 5C**). The DNA origami boxes were stable in FBS in both cations over the timescale of three days. Interestingly, early during exposure to FBS the DNA origami band shifts to higher apparent sizes with both cations (**Figure 4D&H** and **Supplemental Figure 5C**), but this returns to normal values at three days. This may suggest that the hollow DNA origami boxes are either transiently coated by serum components or transiently aggregate.^22,35,37,38^ In contrast, other groups have shown that simpler DNA origami structures are considered to be completely dissembled after 24 h in the presence of 10-20% FBS.^22,35,37^ However, complete disassembly would not explain the DNA origami box shift back to its original band location after three days. As with the other conditions, the hollow DNA origami boxes showed robust stability in serum and displayed no smearing or changes in band intensity. Overall, the DNA origami boxes are outstandingly stable under a variety of conditions that they are expected to encounter during the drug loading process or within the body.

### 2.3 Small Molecule Loading

Once we determined the stability of the hollow DNA origami boxes, we were interested in whether we could load them with a variety of different cargos including both small molecules and large proteins. Around 90% of the drugs sold globally are small molecules. For DNA origami to be a viable vehicle for drug delivery, it must be able to be loaded with small molecules. The easiest way to load molecules is to passively load them during the assembly process. To test whether our hollow DNA origami boxes could be loaded with small molecules, we selected several different fluorescent small molecules as model cargos: fluorescein, calcein, and malachite green. These were chosen to span the spectrum of charges that potential drug cargos might have. The fluorescent small molecules were added to the assembly mixture at 10 mg/mL and assembled with the DNA origami boxes. Negatively charged fluorescein and neutral calcein did not show evidence of loading into the boxes when evaluated by agarose gel electrophoresis (**Figure 5A & B**). Free dye, seen at the bottom of the gel in both the presence and absence of DNA loading stain, confirms their presence. In contrast, malachite green consistently showed evidence of loading as demonstrated by a band in the absence of DNA loading stain (**Figure 5C**). This band is shifted to higher molecular weights likely due to the change in charge imparted on the hollow DNA origami by the positively charged malachite green, decreasing the overall negative charge of the hollow DNA origami. The intensities of the bands of loaded hollow DNA origami boxes differed depending on whether the fluorescent molecule was added during box assembly or after the box was assembled (**Figure 5A&D**). The lower intensity of the band observed when the fluorescent molecule was added after box assembly likely represented fluorescent molecules bound to the outside of the hollow DNA origami box. The higher fluorescence observed in the band of the assembled box indicates that fluorescent dye was indeed entrapped within the hollow DNA origami box. The fluorescence intensity associated with the boxes changed in proportion to the concentration of malachite green present during box assembly, whereas the fluorescence intensity of boxes incubated with malachite green post-assembly saturated at or before 5 mg/L (**Figure 5D** and **Supplementary Figure 6**). The difference between these intensities and the matched boxes assembled in the presence of malachite green increased monotonically with malachite green concentration. These results suggest saturation of binding sites on the outside of the box while loading inside the box continued to increase with increasing malachite green concentration.

**Figure 5:**
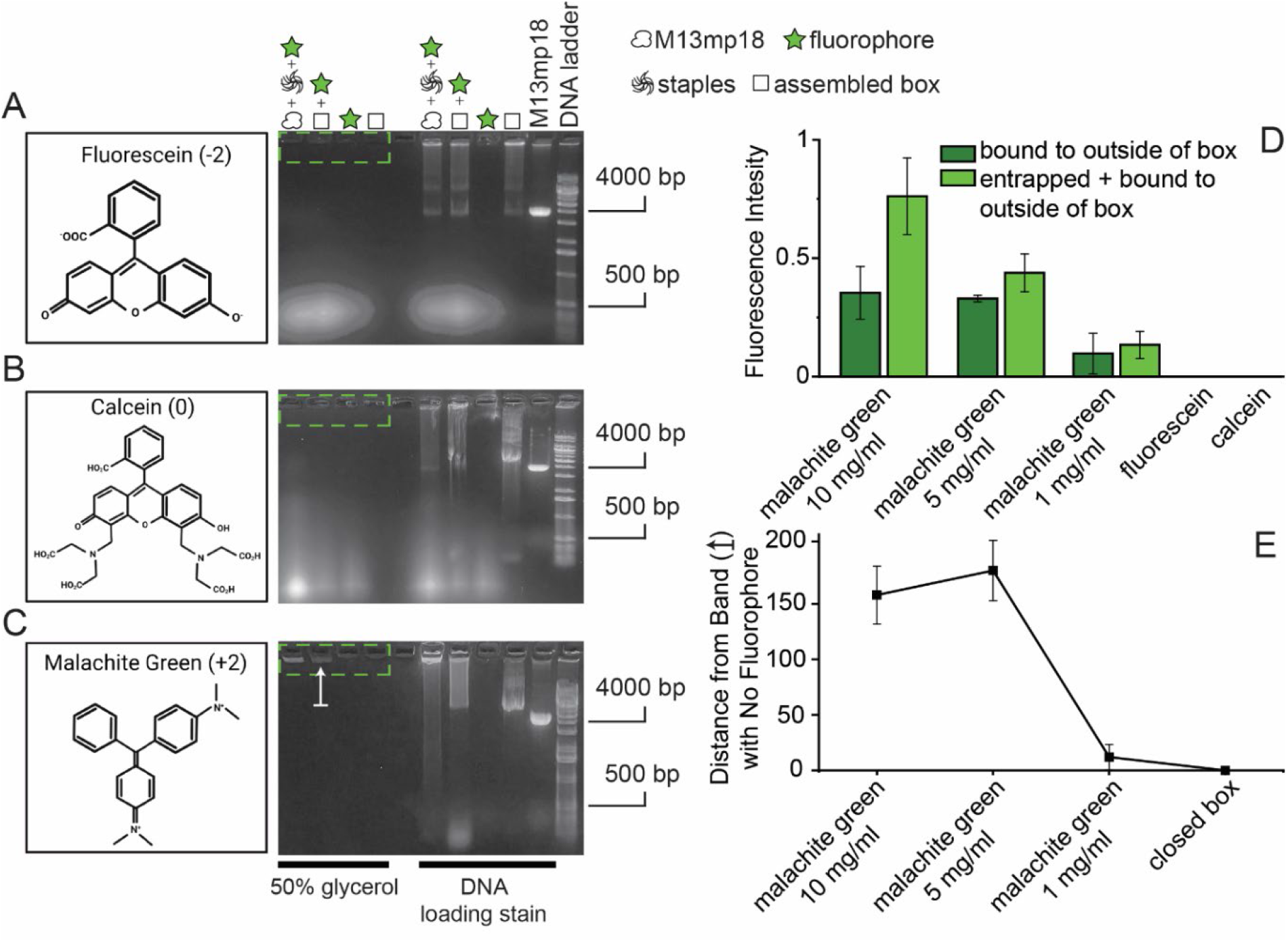
Hollow DNA origami loading is dependent on the charge of the cargo. Fluorescent molecules with A) negative, B) no and C) positive charge were loaded in hollow DNA origami boxes, assembled isothermally. Chemical structure and charge in parentheses (left) and agarose gels of 10 mg/mL fluorescent molecule loaded in 3 µM hollow DNA origami boxes (right) are shown. The dotted green box indicates the location of loaded DNA origami boxes. The white arrow indicates the shift of the band induced by the DNA origami box loaded with fluorophore compared to DNA origami box with no fluorophore. D) Quantification of the band intensity referring to the DNA origami box loaded with fluorophore. Dark green refers to the signal when fluorophore was added to pre-assembled boxes and light green refers to the signal when fluorophore was added during DNA origami assembly. Each bar represents the average of three independent assembly experiments. E) Quantification of the shift of the band induced by the DNA origami box loaded with fluorophore compared to DNA origami box with no fluorophore. Each square represents the average of three independent assembly experiments. All error bars represent 95% confidence intervals.

The shift in box position on the gel is correlated with the externally adsorbed malachite green and not with the presumed trapped malachite green or with the sum of the two. This result is consistent with the assignments of externally adsorbed malachite green, which neutralizes surface charge and internally trapped malachite green, which should have no effect on surface charge (**Figure 5E**). These results demonstrate that positively charged molecules can be passively loaded into hollow DNA origami boxes in a concentration dependent manner.

### 2.4 Protein Loading

#### 2.4.1 Predicting Staple Extension Orientation

Once we were able to load the DNA origami boxes with small molecules, we were interested in whether we could load proteins in the boxes. Passive loading as demonstrated with small molecules requires high concentrations of the small molecule in the assembly mix. This is likely not feasible for proteins due to solubility and expense constraints. Therefore, an active loading process was used, which involved cutting specific staples at a particular position and extending them with biotinylated 3’ or 5’ ends that could capture biotin-binding cargos. With the knowledge that the average helical twist has approximately 10.4 nucleotides, not considering relaxed or supercoiled DNA^39^, we designed four staples, each with starting positions that were shifted three nucleotide along the helix within an eleven nucleotide repeating unit and a biotinylated five nucleotide extensions of the 3’ or 5’ ends (**Figure 6A**). The extensions were exposed to streptavidin labeled beads and the beads were washed to remove unbound material. (**Figure 6B**). DNA origami that attached to the beads was extracted by exposure to NaOH (**Figure 6B**). The NaOH was neutralized with HCl and the two fractions, extract and beads, were resolved by electrophoresis through an agarose gel (**Figure 6C**) and quantified (**Figure 6D**). Boxes with staple extensions on positions 1 and 4 bound to the beads, whereas boxes with staple extensions on positions 2 and 3 remained in the buffer, indicating that staple extensions on positions 1 and 4 were on the outside of the boxes, whereas staple extensions at positions 2 and 3 were on the inside of the box. Using this information, we correctly predicted two more inside staple positions (**Supplemental Figure 7**). Staple extensions can now be added anywhere on the DNA origami box with an accurate prediction of whether they extend into or out of the hollow DNA origami. This capability introduces the possibilities of actively loading protein or of tethering molecules to the outside of the DNA origami.

**Figure 6:**
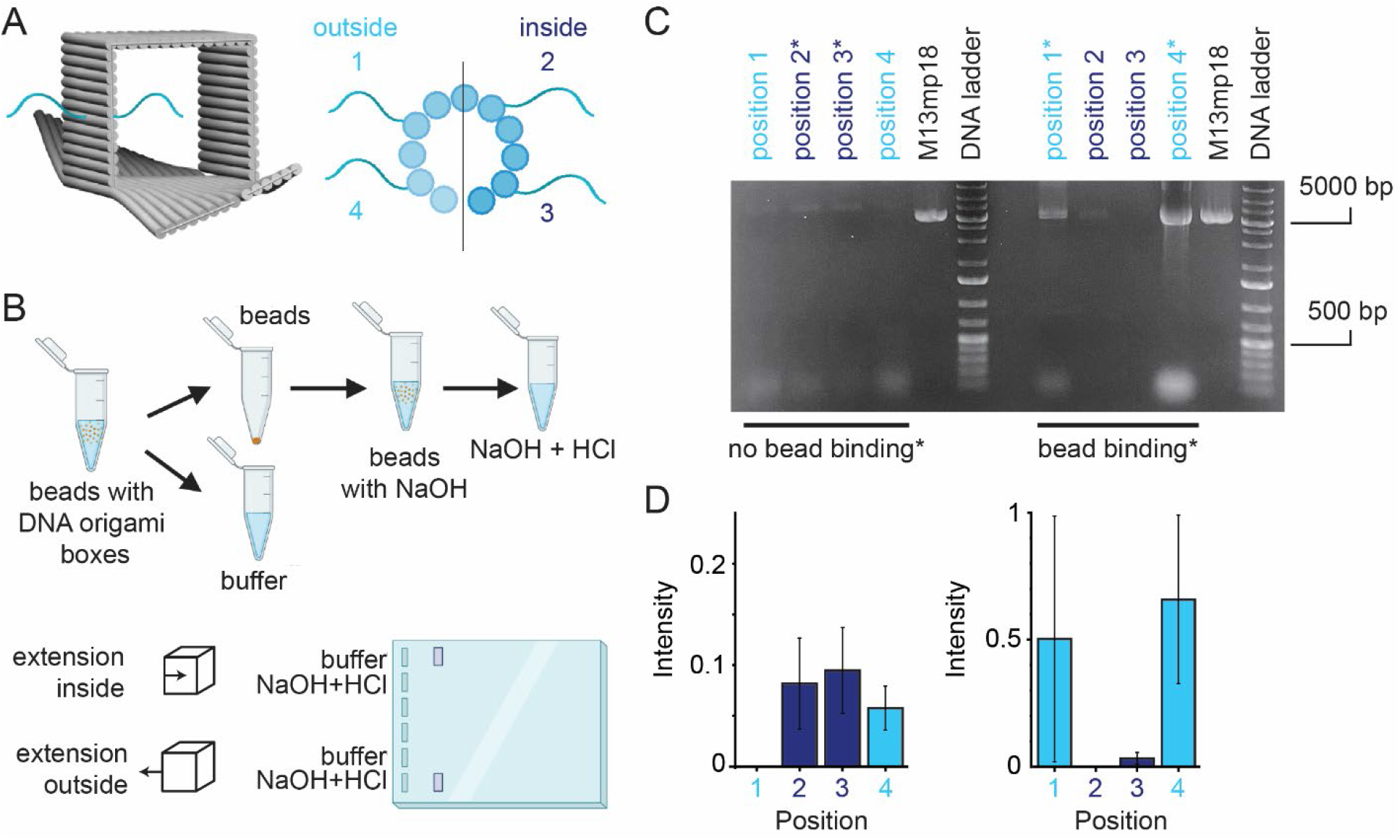
Staple extensions from the hollow DNA origami box can be designed for capture of cargo inside or for decoration with targeting molecules outside. A) Schematic of staples extending inside and outside of the hollow DNA origami box (left). Close up of a single DNA helical twist, showing the direction of extension of different staples. B) Schematic of the experimental process of attaching hollow DNA origami boxes with biotinylated extended staples to streptavidin coated beads to determine if the extension was directed toward the inside or outside. C) Agarose gel showing the binding capabilities of hollow DNA origami boxes with staple extensions at different positions. Light blue refers to staples oriented outside and purple refers to staples oriented inside. The no bead binding wells are samples taken from the supernatant after beads were spun down. The bead binding wells are samples taken from the solution after beads were exposed to NaOH, removing the biotinylated hollow DNA origami boxes from the streptavidin coated beads and subsequently neutralized with HCl. D) Quantification of the band intensity of the hollow DNA origami box. Each bar represents the average of three independent isothermal assembly experiments. All error bars represent 95% confidence intervals.

#### 2.4.2 Protein Loading

Once we determined the orientation of extended staple tethers, we loaded the hollow DNA origami boxes with biologics. Biologics (proteins, antibodies, *etc*.) are used for a number of disease treatments, so a viable way of loading proteins in delivery systems is needed. As mentioned above, passive loading of large molecules is prohibitive due to the challenges of solubility limits and cost. Active loading, using staple extensions that can attach to proteins through an affinity tag provides an effective way to load hollow DNA origami boxes. As a model system we used biotinylated staple extensions and streptavidin. A variety of other nucleotide-protein or functionalized nucleotide tethers including thiol-modified staples, complementary ssDNA strands, zinc “fingers”, antibody fragments and more could be used to actively load protein into DNA origami boxes.^8,26,28,30,40^ Staple extension length likely determines whether a protein can bind to the staple extension. Indeed, others have shown that the length of the tether of surface-attached ligands influences the kinetics of protein attachment, with an optimal tether-length providing for the highest affinities.^41,42^ We designed two staple extensions of five nucleotides (short) and twenty nucleotides (long) (**Figure 7A**). To visualize streptavidin we used FITC-biotin to occupy on average two of the four sites on the protein, which was optimized by titration (**Supplemental Figure 8A**). The remaining sites would be available for binding to the biotinylated staple extensions.

**Figure 7:**
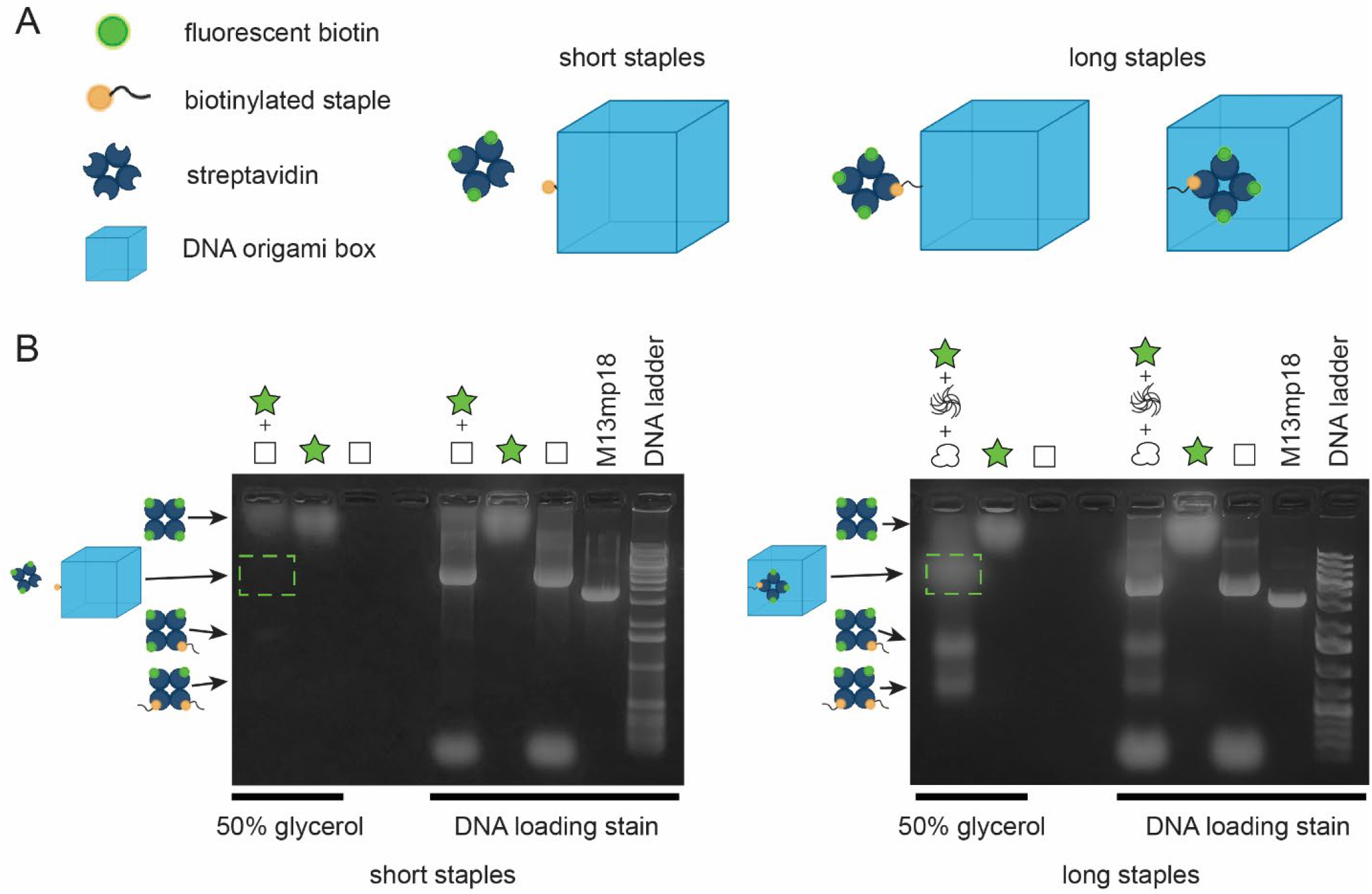
Hollow DNA origami loading requires inside staple extensions of a minimum length. A) Schematic showing that short biotinylated staples on DNA origami boxes, assembled isothermally, do not attach to streptavidin, while long biotinylated staples on DNA origami boxes can attach to streptavidin. Components not shown to scale. B) Agarose gels of DNA origami boxes with short, outside (left) and long, inside (right) extensions. Green dashed box refers to the hollow DNA origami box with attached streptavidin bound to fluorescent biotin. Schematics of other species of bound streptavidin are shown to the left.

We loaded the hollow DNA origami boxes with the streptavidin conjugate by adding the FITC-biotin-streptavidin to the assembly mixture. The assembly mixture is initially heated to 95 °C. Therefore, we considered the temperature sensitivity of the streptavidin-biotin interaction. While streptavidin is only thermostable to 75°C, conjugated with biotin, the thermal stability of the protein conjugate increases to approximately 112 °C.^43^ In addition, we showed that streptavidin heated to 95 °C does not bind to FITC-biotin, but FITC-biotin-streptavidin heated to 95 °C is indistinguishable from non-heated FITC-biotin-streptavidin when run on an agarose gel (**Supplemental Figure 8B**). DNA origami boxes with outside staple extensions of five and twenty nucleotides assembled in the presence of FITC-biotin-streptavidin were resolved through an agarose gel (**Figure 7B and Supplementary Figure 9A**) in the presence or absence of DNA loading stain. A band at the DNA origami box size (green dashed box) was observed in the absence of DNA loading dye for only the twenty-nucleotide staple extension even after 6 h of incubation (**Supplementary Figure 9A**). Similarly, DNA origami boxes with inside staple extensions of five and twenty nucleotides assembled in the presence of FITC-biotin-streptavidin were resolved through an agarose gel (**Supplementary Figure 8A and Figure 7B**) in the presence or absence of DNA loading stain. A band at the DNA origami box size (green dashed box) was observed in the absence of DNA loading dye for only the twenty-nucleotide staple extension even after 6 h of incubation (**Supplementary Figure 8A**). There is likely steric hindrance inhibiting the binding of the FITC-biotin-streptavidin to the DNA origami boxes as the five-nucleotide extension is approximately 1.7 nm in length compared to 5 nm in hydrodynamic diameter of the streptavidin. Biotin binding with streptavidin results in almost an engulfment of the biotin as opposed to an adsorption, explaining biotin’s extremely high affinity binding with streptavidin.^44^ When the staple is extended to 6.6 nm, the streptavidin can more easily attach to the biotinylated staple both inside and outside of the DNA origami boxes. These binding experiments indicate that staple extensions of longer than five nucleotides are needed to either bind targeting proteins on the outside of the hollow DNA origami box or to actively load protein cargo into the cavity of the hollow DNA origami box. Taken together these experiments outline the design principles associated with tethering targeting molecules to the outside of a hollow DNA origami box and actively loading it with protein cargo.

## 3. Conclusions

In this paper, we report the design and assembly of a unique hollow DNA origami structure with multiple lids. We have characterized the sizes and shapes of these structures using a variety of techniques including in-solution AFM. We have developed an approach to isothermally assemble hollow DNA origami structures across a variety of temperatures over the timescale of minutes, providing a means of rapidly assembling many boxes. While these hollow DNA origami structures appear to be sensitive to centrifugal force, they are exceedingly stable at different temperatures, pH and in different salts as well as in the presence of both solvents and biologically relevant fluids. We demonstrated that these hollow DNA origami boxes are agnostic to cargo size, loading both small molecules and proteins. However, the charge of the small molecule governs the ability to passively load hollow DNA origami boxes origami boxes.

Active loading of large proteins can be facilitated by extending staples into the hollow DNA origami box cavity, however staple extension length must be optimized to enhance affinity and to overcome steric hindrance. The internal tethers could be temperature labile, binding cargo at cold temperatures during assembly with release under higher temperatures after the DNA origami box is closed, priming the cargo for release upon lid opening. These DNA origami boxes with multiple lids have the potential to be used with locking mechanisms that can be opened via different inputs to release a variety of cargos in drug delivery applications.

## 4. Materials and Methods

### 4.1 DNA Origami Design

The design of staple strands to assemble the hollow DNA origami box was done using cadnano square. The predicted 3D structure was created using CanDo and UCSF Chimera imaging.^45–47^ M13mp18 circular bacteriophage DNA was used as the scaffold for staple design. The full staple set is shown in **Table S1**.

### 4.2 DNA Origami Assembly

Hollow DNA origami boxes were assembled in a TAE (40 mM <u>T</u>ris base, 20 mM glacial <u>A</u>cetic acid and 0.8 mM <u>E</u>DTA) buffer with 12.5 mM Mg(OAc)_2_ or 100 mM NaCl. M13mp18 circular DNA (P- 107, Bayou Biolabs) was used as the scaffold for the staples (IDT). Scaffold at 1 to 3 µM with between a 1:2 to 1:5 scaffold:staple molar ratio. For gradient assembly, the assembly solution was split into 50 µL aliquots and placed into a thermocycler to be heated at 95 °C for 5 min and cooled at -1 °C/min until 25 °C. The aliquots were combined after cooling. For isothermal assembly, the assembly solution was heated in a dry bath at 95 °C for 10 min, and then cooled at 4-90 °C for 30-120 min.

### 4.3 DNA Agarose Gel Electrophoresis

Agarose gels (1%) were run at 5.3 V/cm for 60 min. Prestain or loading stain was used to visualize the DNA bands. If prestain was used 6 µL of prestain (SmartGlow Prestain, E4500-PS, Southern Labware) was added to 60 mL of agarose gel. If loading stain was used, an appropriate volume of 6x loading stain (SmartGlow Loading stain, E4500-LD, Southern Labware) was added to bring the sample to 1X loading stain before loading into the gel. For fluorescent cargo experiments DNA stains were not used and in place of 6x loading stain, a 50% (v/v) glycerol-in-water solution was added to the sample at the same volume as the loading stain. Gels were analyzed using the Gel Analyzer tool in ImageJ. The analysis area for each gel extended from immediately after the well to immediately after the staple bands. The intensities of the bands were determined by finding the area under each peak of the averaged linescan, which accounts for background fluorescence calculated locally.

### 4.4 DNA Origami Purification

Purification was done using Amicon 100k and 50k MWCO filters (UFC5100, Sigma). The manufacturer recommends that the filters be spun in the centrifuge at approximately a centrifugal force of 20k x *g* for 5 min, which was later lowered to 5k x *g* for 20 min. Alternatively, the following procedure for PEG purification was done for loading experiments to prevent clogging of the filters and to limit expense. Large amounts of dye used in loading experiments required several filter passes. This either resulted in clogged filters if the same filter was used or the necessity to use multiple filters. 50 µL of PEG (17 wt% 8,000 Da with TAE and 12.5 mM Mg(OAc)_2_) was added to 400 µL of assembled DNA origami. The solution was gently mixed and place at 4 ⁰C for 10 min before being spun at 12.6k x *g* for 30 min.

The top 400 µL was removed and 350 µL of TAE with 12.5 mM Mg(OAc)_2_ was added along with another 50 µL of PEG solution. This was repeated until the visible dye was dramatically diluted (1-4 repetitions).^17^

### 4.5 Dynamic Light Scattering (DLS)

DLS was performed using the Zetasizer Nano-ZS (Malvern). With ZEN0040 equivalent cuvettes, 50 μL of either filter purified from staples or unpurified DNA origami boxes was used determine the size. Water was selected as the dispersant and a refractive index of 1.53 was used for the DNA origami size determination. The system temperature was set to 25 °C with an equilibration time of 120 s for data in **Figure 1**. For isothermal data in **Supplementary Figure 3**, the system temperature was set to 4°C.

### 4.6 Atomic Force Microscopy

#### 4.6.1. Sample Preparation

A muscovite mica (grade V1) substrate was subjected to a fresh cleaving process to achieve optimal surface conditions for Atomic Force Microscopy (AFM) imaging. Subsequently, a 200 μL volume of a purified DNA origami sample was combined with 100 μL of a TAE buffer containing 12.5 mM Mg(OAc)_2_. The resultant mixture was gently deposited onto the surface of the mica substrate.

Following deposition, adsorption was allowed to proceed for 35 min at room temperature before initiating the AFM experiment.

#### 4.6.2. Instrumentation

AFM imaging was conducted utilizing a BioScope Resolve system (Bruker), equipped with a SCANASYST-FLUID probe featuring a spring constant of 0.4 N/m and tip radius 20 nm. Imaging was executed in ScanAsyst mode. The BioScope Resolve AFM was sitting inside a vibration and noise isolation chamber that reduces noise level in nanoscale measurements. It was also sitting on top of a vibration isolation table, which needed to be switched on prior to each experiment. There was a magnetic contact on the base plate/sample stage used to lock the stainless steel discs that were attached to the mica used for sample imaging. All imaging procedures occurred in solution within a controlled environmental chamber, which was isolated to minimize potential sources of drift and temperature fluctuations.

#### 4.6.3. Imaging Parameters

AFM images were acquired at a variable scan rate spanning between 0.3 and 0.4 Hz to prevent damage to the DNA origami sample and to yield topographical and structural details. The scan size was modulated within the range of 3-7 μm to capture diverse regions of interest on the muscovite mica substrate and the AFM height images were zoomed in on locations with significant features. A consistent peak force set point of 200 pN was maintained to preserve the structural integrity of the DNA origami.

Imaging utilized a resolution of 256 pixels per line. The peak force amplitude setpoint and peak force frequency were kept constant at 250 nm and at 1 kHz respectively during imaging.

#### 4.6.4. Data Analysis

Nanoscope Analysis and WSxM 5.0 software were employed to process and analyze the AFM images acquired during the experiment. Structural attributes, including dimensions and topographical features of the DNA origami structures, were extracted from these images. Shape characteristic data from was found by manually counting the number of distinct shapes across four images for two independent assemblies per DNA origami box design. Purification data was quantified by manually counting the number of small molecules (size less than ∼25 nm) and DNA origami boxes (size approximately 30-45 nm) in one image per condition.

### 4.7 Stability

Stability at different temperatures was tested as follows with a base case of 4 °C, pH 7.5, with 12.5 mM Mg(OAc)_2_ and no DMSO or FBS. For example, when temperature was changed, pH and ion remained the same as the base case. Unpurified DNA origami closed boxes were kept at 4 °C, 18 °C and 37 °C for up to 14 days. Equal volumes of each sample from each timepoint for each temperature were loaded into the gel (approximately 10 µL/sample). Stability as a function of pH was tested as follows. Filter-purified DNA origami closed boxes were kept at pH 7.5 and pH 5 for up to 14 days. Both buffers were TAE with 12.5 mM Mg(OAc)_2_. The pH 5 buffer was adjusted using acetic acid and the pH 7.5 buffer was not adjusted. DNA concentrations of filter-purified DNA origami boxes were measured using a spectrophotometer to check for stability based on mass. The pH 5 samples were at a 4:1 volumetric ratio of pH 5 TAE buffer to filter-purified DNA origami boxes in pH 7.5 TAE. Equal mass of DNA origami was loaded into the gels for each timepoint for each condition. Stability in DMSO and FBS was tested as follows. Unpurified DNA origami closed boxes were kept in 10% DMSO or 10% FBS (Non-heat inactivated, A5209401, Gibco) for up to 3 days. In experiments using DMSO, the DNA origami closed boxes were assembled with 12.5 mM Mg(OAc)_2_. In experiments using FBS, the DNA origami closed boxes were assembled with 12.5 mM Mg(OAc)_2_ or with 100 mM NaCl. Equal volumes from each stability sample from each timepoint for each condition were loaded into the gel (approximately 10 µL/sample).

### 4.8 Loading hollow DNA origami boxes with fluorescent small molecules

DNA origami boxes were loaded with molecules of different charge: fluorescein (-2) (863200, Carolina Biological Supply), calcein (0) (C0875, Sigma), or malachite green (+2) (32745, Sigma). Closed DNA origami (3 µM) boxes were either assembled with fluorescent molecule to measure how much was entrapped and bound to the DNA structure or after assembly to measure how much was bound to the DNA structure. Final concentrations of fluorescent molecules present during assembly ranged from 1-10 mg/mL. Loaded DNA origami boxes were purified using up to four rounds of PEG purification to reduce background fluorescence. Some loaded DNA origami samples showed a mobility shift due to a difference in charge between the loaded DNA origami structure and DNA origami structure with no dye. Under some conditions, the DNA origami structure moved more slowly than the largest ladder band (approx. 20 kb). These structures were quantified in pixels rather than by estimates of apparent size. Gel pictures were taken at a consistent height using the SmartDoc Gel Imaging System (Accuris Instruments) with Blue Light Illumination.

### 4.9 Extending DNA staples

Extended DNA staples were designed by cutting the original staple at a designed position and extending it with a 3’ or 5’ poly-A or poly-T tail of lengths 5 or 20 nt long. The remaining small fragment was not used in the assembly. The cut was made to minimize the length of original sequence that was not used in the assembly. The unused sequence was no more than 10 nt long. The polyA or polyT segments were not complementary to the scaffold. Once a position was identified as either inward facing or outward facing, one could determine other inward or outward facing positions along a single helical twist by counting 11 nt in the cadnano design, based on the assumption of normal tension and pitch. The extended staples were biotinylated on the 3’ or 5’ end of the extensions that would be inside or outside of the DNA origami boxes. All staples extended for Figure 6 were biotinylated at the 3’ end. All staples extended for Supplementary Figure 7 were biotinylated at the 5’ end. Figure 6 position 1 staple extension 1: 5’-TGC CGT CGA TTT CGG AAC CTA TTA AAA A-3’. Figure 6 position 2 staple extension 1: 5’- TGC CGT CGA TTT CGG AAC CTA TTA TTA AAA A-3’. Figure 6 position 3 staple extension 1: 5;- TGC CGT CGA TTT CGG AAC CTA TTA TTC TAA AAA-3’. Figure 6 position 4 staple extension 1: 5’- TGC CGT CGA TTT CGG AAC CTA TTA TTC TGA AAA AA-3’. Supplementary Figure 7 (B) staple extension: 5’-TTT TTA GAG CTC CAA CGT CAA CCT CAG AAC CGC CAC CC-3’. Supplementary Figure 7 (C) staple extension: 5’- TTT TTA CAT GGC TTT TGA TGA TAC AGG ATT GGC CTT-3’. For experiments identifying inward or outward facing extensions, the DNA origami boxes were exposed to streptavidin-coated beads (MyOne Dynabeads, 65001, Thermofisher), gently mixed for 10 min and the supernatant (no bead binding supernatant) was collected after pulling down the beads with a magnet. Then the beads were exposed to 10 µL of 100 mM NaOH, gently mixed for 10 min and the supernatant (bead binding supernatant) was collected and added to 10 µL 100 mM HCl to neutralize the solution. No bead binding and bead binding supernatants were loaded into agarose gels and analyzed as described above.

### 4.10 Loading hollow DNA origami boxes with fluorescently-labeled protein

There were short poly-A extensions of five nucleotides and long poly-A extensions of twenty nucleotides from the original staple extensions that were directed inward or outward of the DNA origami boxes (listed above). The extended staples were biotinylated on the 3’ end of the extensions that would be inside or outside of the DNA origami boxes. This would allow them to bind to streptavidin (21122, Pierce). Streptavidin was bound with fluorescein-isothiocyanate (FITC)-biotin (30574-1mg-f, Atto) (1:2 molar ratio of streptavidin:FITC biotin). Streptavidin-FITC-biotin conjugate was added at a 1:1 molar ratio to the biotinylated extended staples. When using the short, outside extended staples, DNA origami boxes with the staples were exposed to the FITC-biotin-streptavidin conjugate for approximately 6 h.

When using the long, outside extended staples, DNA origami boxes with the staples were exposed to the FITC-biotin-streptavidin conjugate for approximately 30 min. When using the short or long inside extended staples, isothermal assembly of the DNA origami boxes at 4 °C for 30 min was done with the FITC-biotin-streptavidin conjugate.

## Acknowledgements

We thank Lee Bendickson for his technical guidance on using DNA origami design tools as well as the rest of the DNA Origami working group at ISU for their helpful input. **Figures 6, 7**, S8, and S9 were created with Biorender.com.

## Abbreviations

DNA: 
DLS: 
AFM: 
DMSO: 
NaCl: 
FBS: 
Mg(OAc)_2_: 
TAE: 
IDT: 
PEG: 
FITC: 

## Author Contributions

A.H.K.: Conceived and conducted the experiments, analyzed the data and interpreted the data across all figures, wrote the article, prepared the figures and edited the article. N.M..: Conducted the experiments and interpreted the data in figures 1, 3 and S2 and edited the article. J.V.H.: Conducted the experiments, analyzed the data and interpreted the data in figures 4, S3 and S4. C.S.: Conducted the experiments in figures 1, 3 and S2. M.N.H: Participated in discussions of the data, their interpretations, made suggestions for experimental design, and edited the article. A.S.: Conceived and conducted the experiments and interpreted the data in figures 1, 3 and S2 and edited the article. I.C.S.: Conceived the experiments, analyzed the data and interpreted the data across all figures, wrote the article, prepared the figures and edited the article.

