## Supplementary Information for "Design and Assembly of a Cargo-agnostic Hollow Two-lidded DNA Origami Box for Drug Delivery"

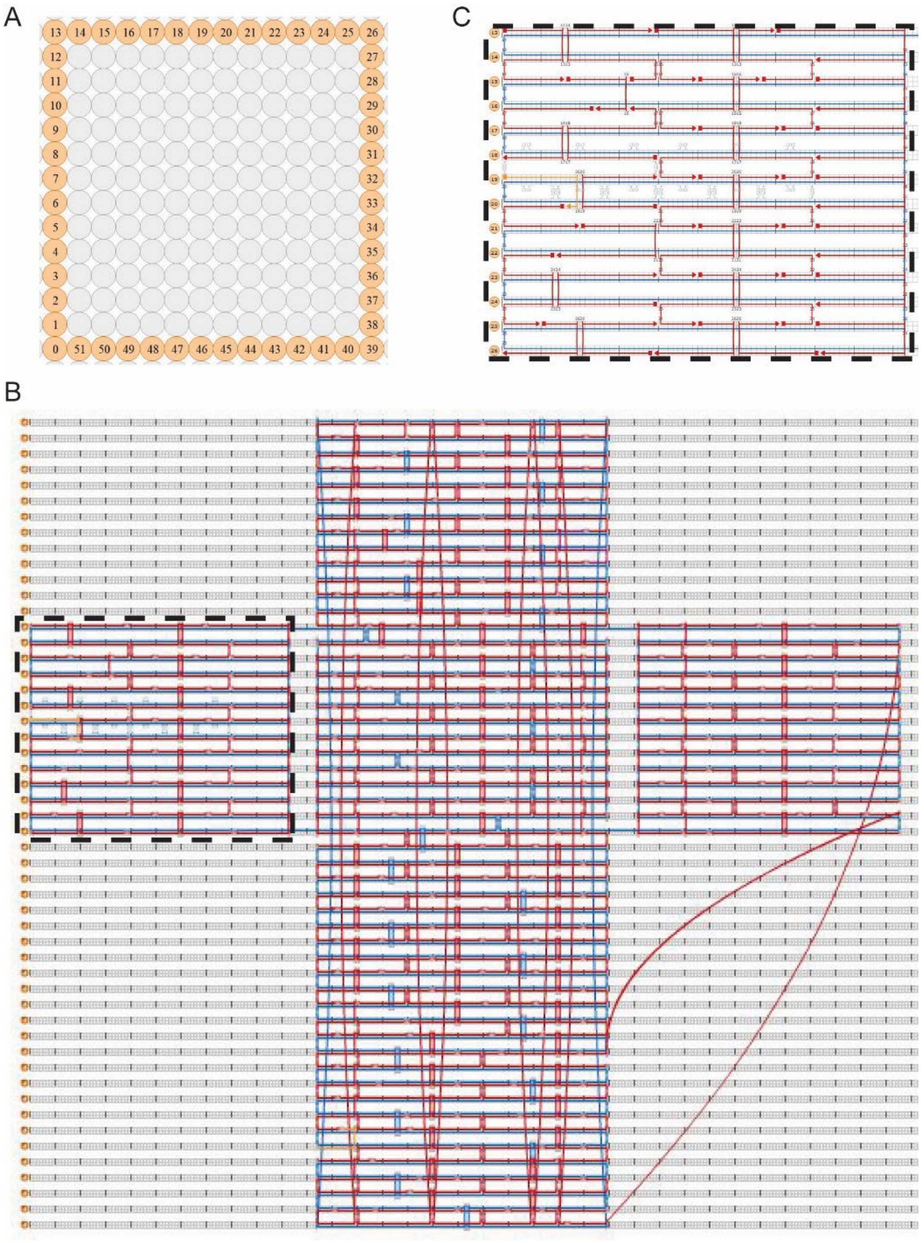


**Supplementary Figure 1: Design of the hollow DNA origami structure.** A) Layout of the DNA helices in the cadnano square view. B) cadnano square 2D design of the one lid open hollow DNA origami box. The region that was enlarged is shown in dashed lines. C) Enlarged region of one lid open design with the extended staple for the lock is in the middle of the left side and is highlighted yellow.

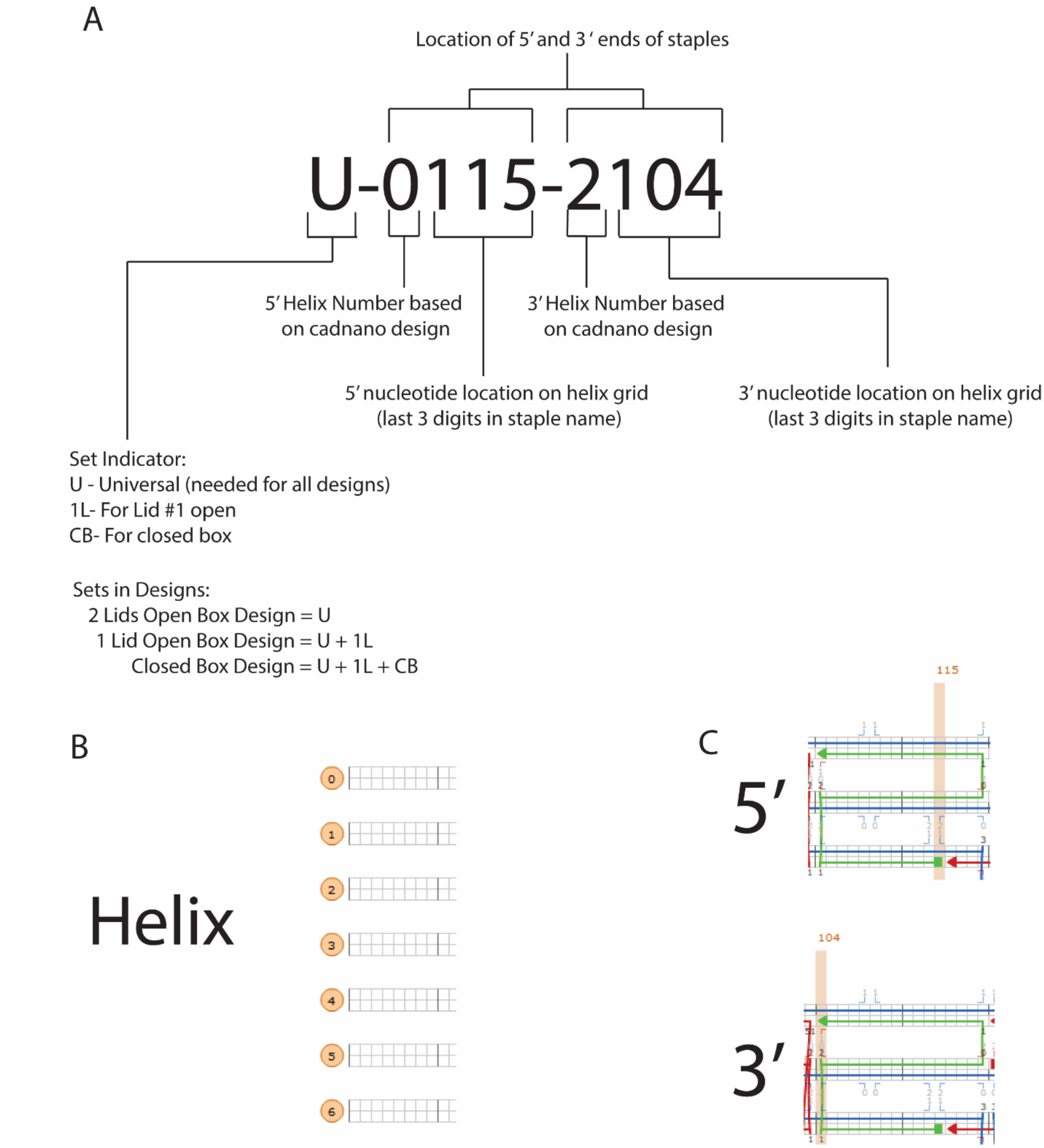


**Supplementary Figure 2:** Staple Name Guide A) The name guide based on specific designs, helix, and 5’ and 3’ locations. B) How to determine helix location. C) How to determine the 5’ (squares) and 3’ (arrow heads) locations.

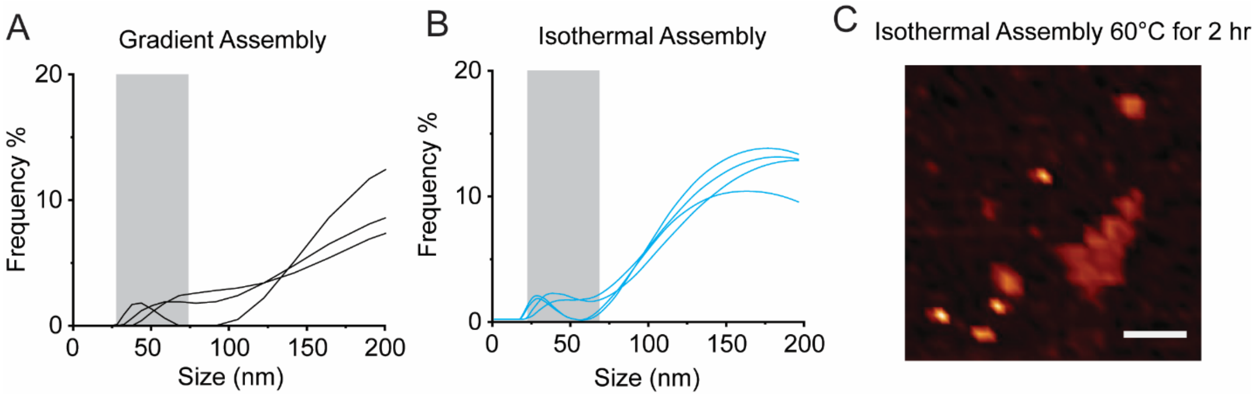


**Supplementary Figure 3: Isothermal assembly of hollow DNA origami boxes results in similar sizes.** A) Size distribution from DLS of unpurified hollow DNA origami closed boxes assembled using a temperature gradient. B) Size distribution from DLS of unpurified hollow DNA origami closed boxes assembled using isothermal assembly at 0 °C for 10 min. Grey boxes refer to size ranges predicted for DNA origami boxes. C) AFM image of DNA origami closed boxes assembled isothermally at 60 °C for 2 h and filtered purified. Scale bar is 200 nm.

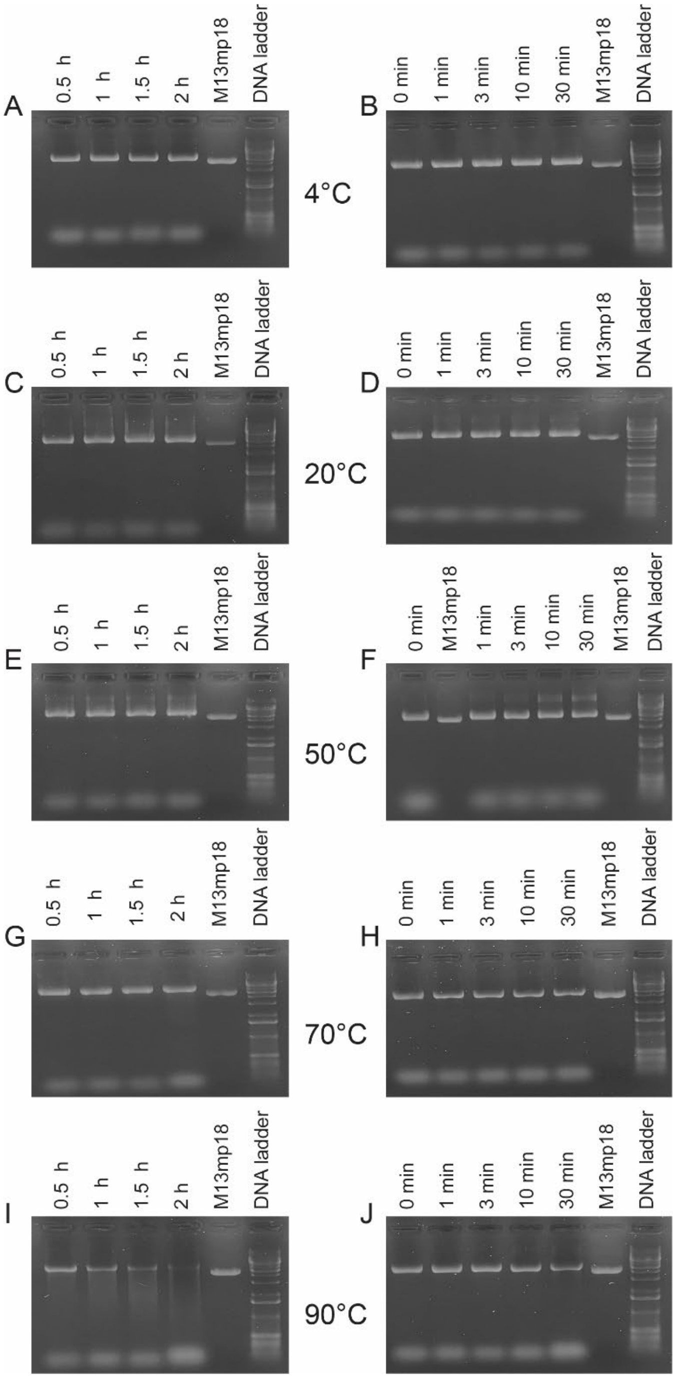


**Supplementary Figure 4: Isothermal assembly of hollow DNA origami boxes occurs rapidly at various temperatures.** Agarose gels of DNA origami closed box isothermal assembly at A & B) 4 °C, C & D) 20 °C, E & F) 50°C, G & H) 70°C and I & J) 90°C from 1 min to 2 h of incubation.

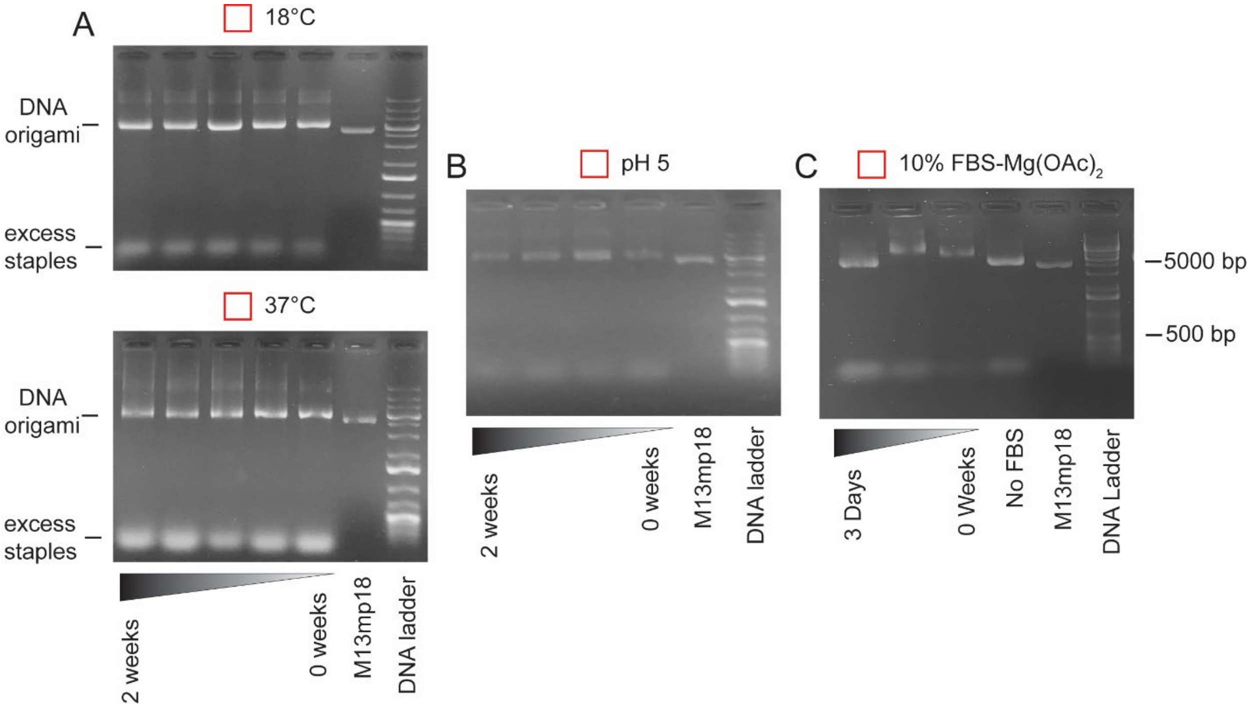


**Supplementary Figure 5: Hollow DNA origami is relatively stable.** Agarose gels of DNA origami closed box isothermal assembly at 4 °C exposed to A) temperatures of 18 °C and 37 °C over the course of 2 weeks, B) pH 5 over the course of 2 weeks and C) 12.5 mM $Mg(OAc)_2$ in 10% FBS over 3 days.

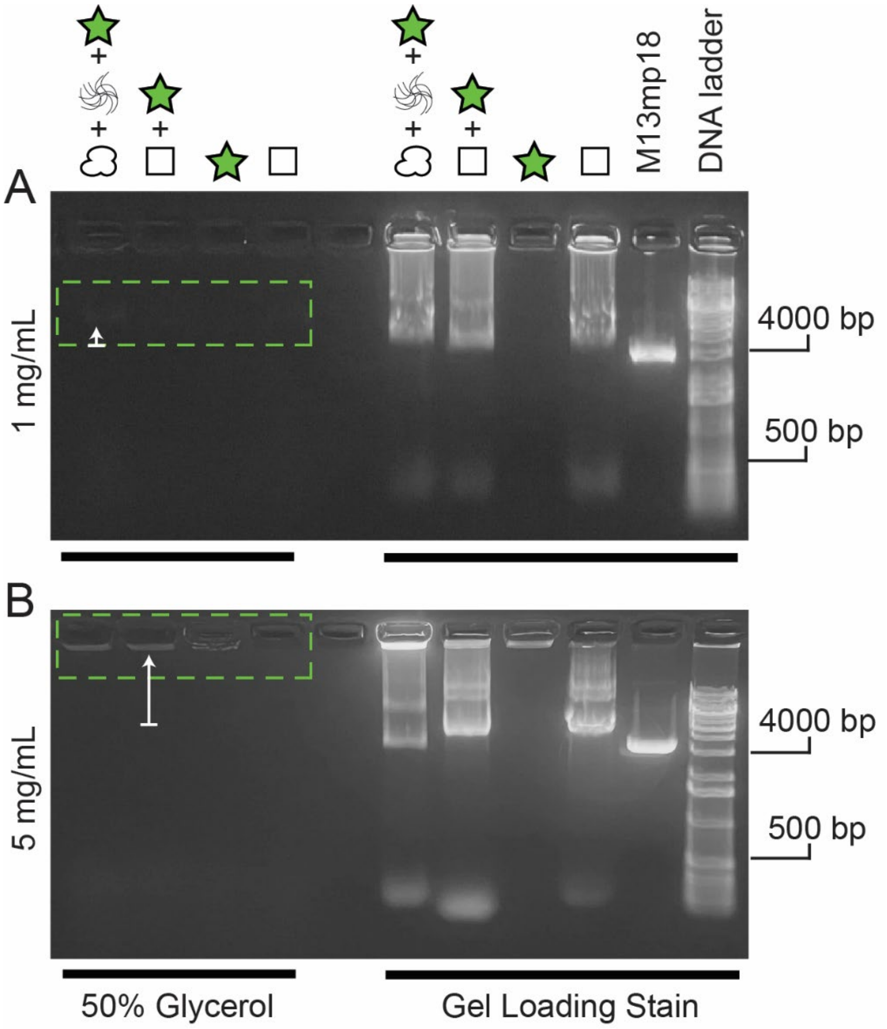


**Supplementary Figure 6: Dose response of malachite green loading in hollow DNA origami boxes.** Agarose gels of A) 1 mg/ml and B) 5 mg/ml malachite green loaded into DNA origami boxes using isothermal assembly. The dotted green box indicates location of loaded DNA origami boxes. The white arrow indicates the shift of the band induced by the DNA origami box loaded with fluorophore compared to DNA origami box with no fluorophore.

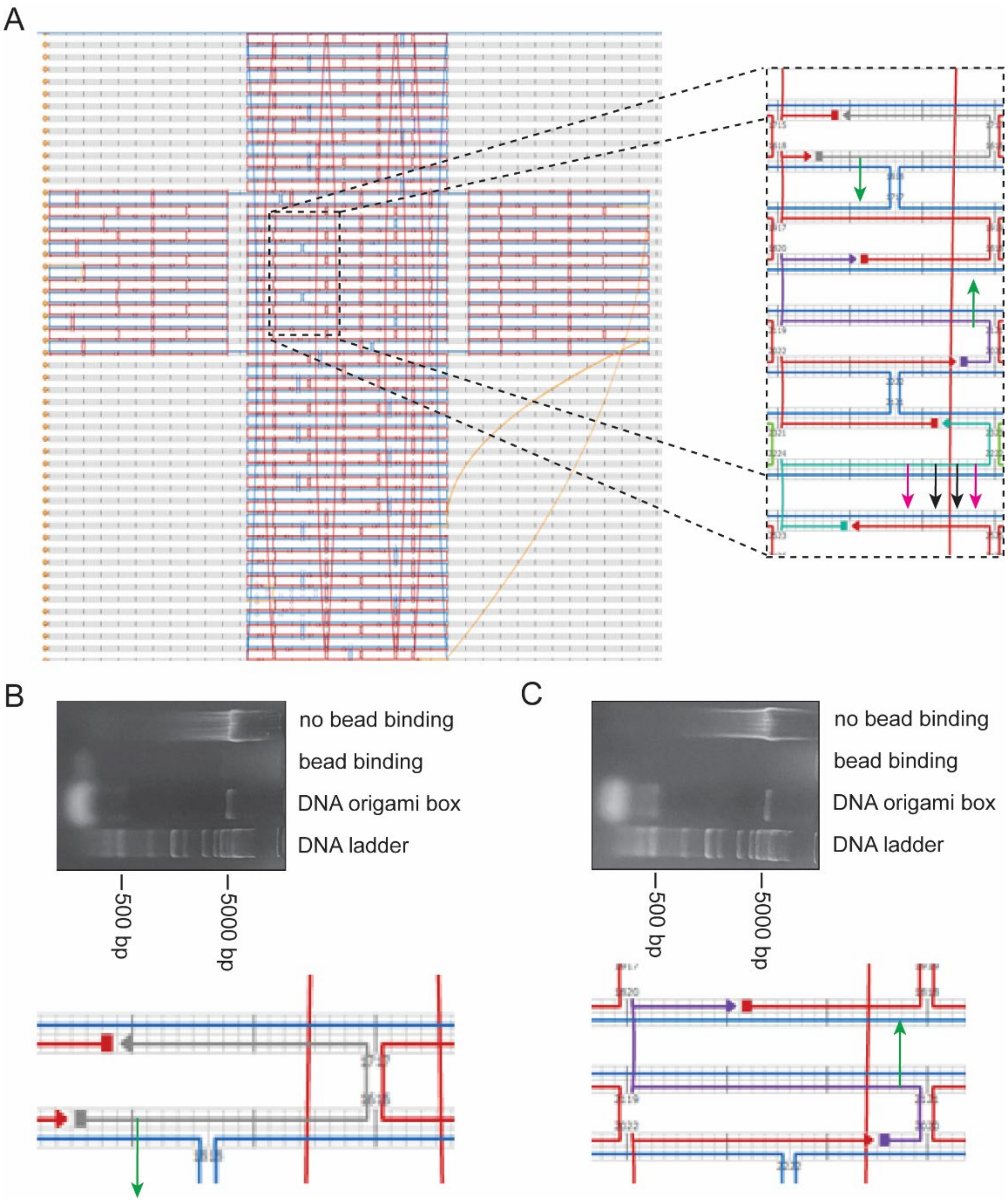


**Supplementary Figure 7: Demonstration of the ability to predict inside vs. outside positioning of staple extension based on helical twist pitch.** A) Cadnano square 2D design of the closed hollow DNA origami box design. The region that was enlarged is shown in black dashed lines. Pink arrows represent staples that primarily extend outside the DNA origami box. Black arrows represent staples that primarily extend inside the DNA origami box (**Figure 6**). Green arrows represent staples extensions that are predicted and confirmed to be primarily inside the box based on the scaffold orientation (above or below the staple strand in the helical grid) and nucleotide location relative to the initial staple extensions from the first experiment. B,C)

Gels that confirm the predicted staple extensions are inside DNA origami boxes. The cathode is on the right, so bands run from right to left.

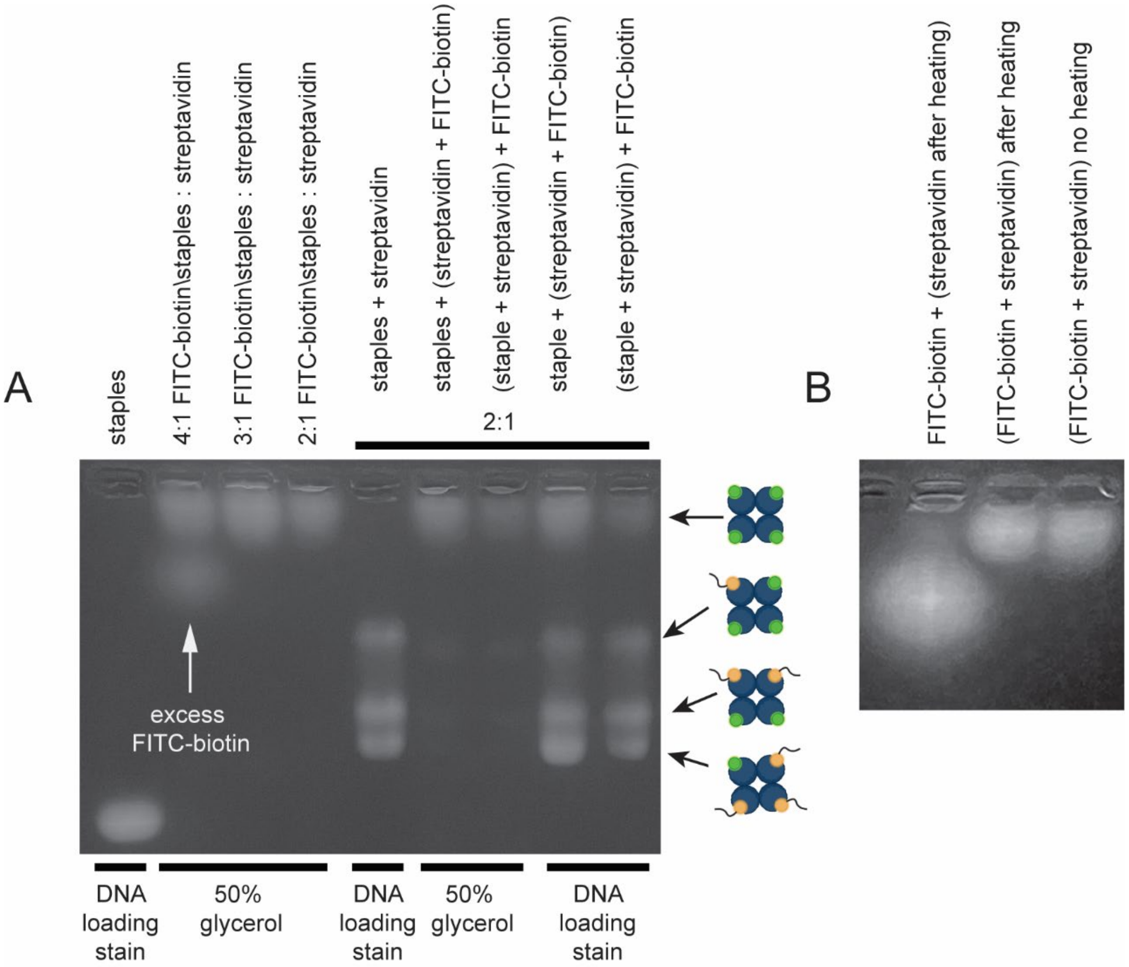


**Supplementary Figure 8: Determination of staples, FITC-biotin, and streptavidin interactions and streptavidin-biotin conjugate thermal stability.** A) A gel showing how staples, FITC-biotin and streptavidin interact. There is a lane containing staples and no streptavidin as reference for other lanes with staples. When FITC-biotin was added to streptavidin at a 4:1 ratio, excess biotin is observed which is not present at 3:1 and 2:1 ratios. When biotinylated staples were added to streptavidin, the observed fluorescent bands represent a distribution streptavidins with varying numbers of staples per tetramer (1, 2, or 3 staples/streptavidin). The lanes on the right side of the gel compare the fluorescent intensity observed after adding the staples first or the FITC-biotin first to the streptavidin (observed with and without fluorescent DNA loading stain). B) A gel showing the thermal stability (95°C) of streptavidin with and without FITC-biotin compared to streptavidin + FITC-biotin not exposed to heat.

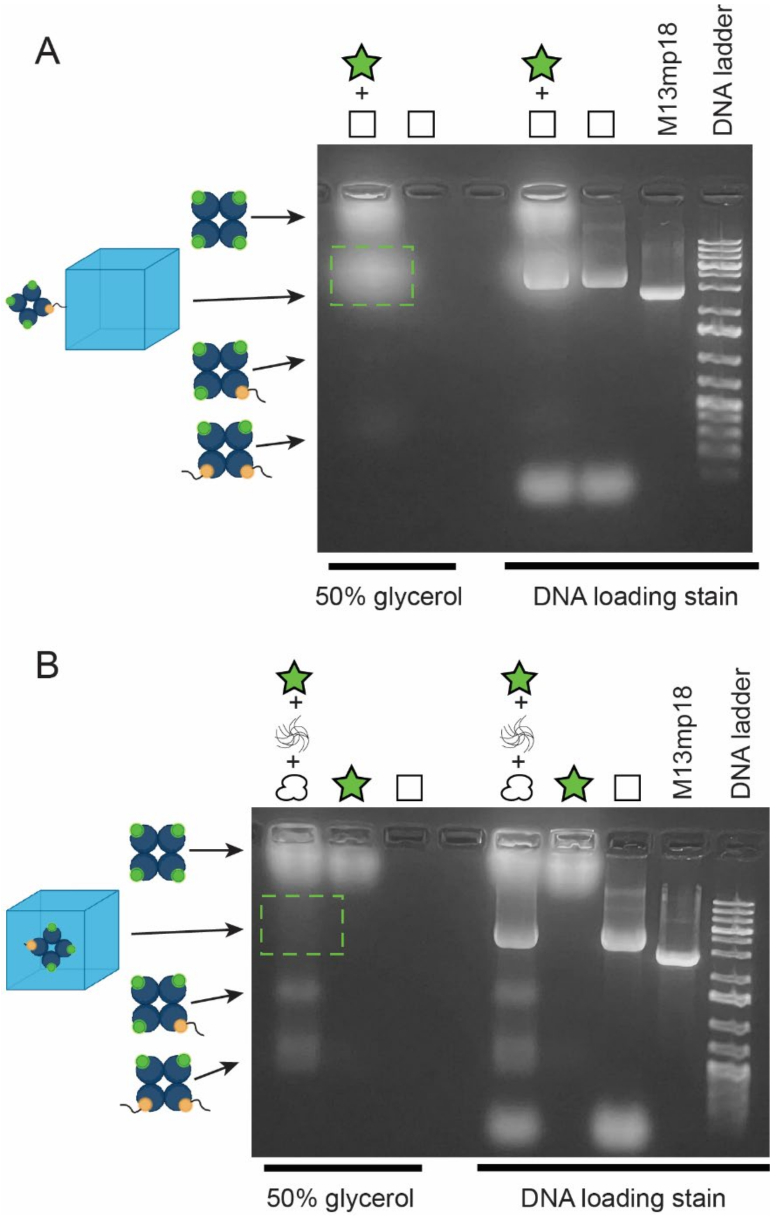


**Supplementary Figure 9: Hollow DNA origami loading requires inside staple extensions of a minimum length.** Agarose gels of DNA origami boxes with A) long, outside and B) short, inside extensions. Green dashed box refers to the hollow DNA origami box with attached streptavidin bound to fluorescent biotin. Schematics of other species of bound streptavidin are shown to the left.

**Supplementary Table 1:** Staple sets for all designs. See Staple Name Guide (Fig. S2A) for specific staple locations.

| Sequence Name | Sequence |
|---|---|
| **U-0143-2144** | CATCACCTGCAACAGTGCCACGCTTAATAGTA |
| **U-1098-3097** | ATAAAAGAACGAACCACCATAATGCG |
| **U-1120-50136** | AGTATTAACACCGCCTTGCTGAACCTAAAATATCTACGCCACATATTA |
| **U-2115-0104** | ATTAAAAATACCCAGAGGTGAGGCGGTCTCAAACCCTCAATCAA |
| **U-2143-4144** | AAATGTTTCATAAATATTCATTGATCTGACCT |
| **U-2158-50163** | GAGGGGGGAGAGCCAGCAATAAACGACGGCACATAGTAAATACC |
| **U-3098-5097** | CGAACTGGCTATTAGTCTTAAATGGA |
| **U-3111-2116** | CTAAAACATAATACTGCGGAATCGTAGACTGGATAGCGTCCCGCC |
| **U-3177-1176** | TCAGAAAGGCTTTTGCAAAAGAAAACCAAAAT |
| **U-4143-1144** | GAAAGCGTAATAAAAGGGACATTCAAAAGATT |
| **U-4158-2159** | GAACCCTATCCCCCTCAAATGCTGTTTTGCCA |
| **U-5098-7097** | TTATTTGCTCAATCGTCTGCCATTGC |
| **U-5129-3110** | GACCAGTAAGAATACGTGGCACAGACAATATTTTTGAATGATAGCC |
| **U-5177-3176** | CCCTGACACGAGAATGACCATAATTAAACAGT |
| **U-6114-5128** | TACATTTTGACACATTGGCAGATTCACCAGTCACAC |
| **U-6143-8144** | AAGAGGAACGAGCTTCAAAGCGAATGAGTAGA |
| **U-6158-4159** | TGCATCATGGCCAACAGAAAATCATCAAGATA |
| **U-7098-9097** | AACAGGATATTACCGCCAGGGCCACC |
| **U-7104-9119** | AAAAACGCTTCCAGAACAAAGAGTCTGTCCATCA |
| **U-7117-6115** | GAAATATCGCGTTTTAATTGCCCGAAAGACTTCAAATACC |
| **U-7177-5176** | GTCAGGATATTATAGTCAGAAGCAGGTCTTTA |
| **U-8143-10144** | AGAACTCATAGCAATACTTCTTTGATAATGCT |
| **U-8158-6159** | ACTTGCCCCAGACCGGAAGCAAAAAAGCGGAT |
| **U-9098-11097** | GAGTAATTATAATCAGTGAGGGATTT |
| **U-9120-11114** | CGCATATGATCCTGAGAAGTGTTTGGAACGGTACG |
| **U-9132-7116** | GTTGAACTATCGGCCTTGCTGGTAATACATG |
| **U-9177-7176** | GGTCATTTTAGAGAGTACCTTTACTCCAACAG |
| **U-10143-12145** | GTAGCTCATCTGGAAGTTTCATTCCACGTAT |
| **U-10161-8159** | ATTGCTGAATATTAGTAATAATCCTTATTGCCATC |
| **U-11098-13103** | TAGACAGAGGCCGATTAAATTAGCGTCAGACT |
| **U-11115-13107** | CCAGACAACTCAGAGCGGGAGCTAAACAGGTAC |
| **U-11130-9131** | ACGGTGACATGTTTTAAAAATTAACC |
| **U-11168-9176** | CCCAATTCTGCGAACGTTTGCGGATGGCTTAGTTGATAAGA |
| **U-12144-14136** | AACGTGCTTCGCCGCTAGCGCTAGG |
| **U-12165-10162** | ATTGCTTTGACGAGCATATAACAGTTGATTAGCTTA |
| **U-13031-15039** | TCACCAATGAAACCATCGTCACCGACTTGAGCCCCAAAGAC |
| **U-13056-14064** | CACCGTAATCAGTAGCGACAGAATCAATGACGGAAATTATTCATTA |
| **U-13108-14091** | GTAACTGCGGCGCGTTTTCATCGGC |

| | |
|---|---|
| **U-13122-11129** | CGCCGCGCTTAATGTCCTCGTTAGAATAAAGT |
| **U-13232-15232** | GGCATCAAAAAGCCTCAGAGCATAATTTCAACG |
| **U-13249-13276** | TAGTAGTAGCATTAACATCCAATAAATC |
| **U-14135-16136** | GCGCTGGCTTTTCATAATCAAAATCCCTCAGA |
| **U-14151-14168** | AGGAGCGGCAGGGCGCGTACTATGGTTAGATACATTTCGCGAGAAAGG |
| **U-14167-16168** | AAGGGAAGAGAGCTTGACGGGGAAAGGGAGCC |
| **U-14183-12166** | AACGTGGCAAATGGTCAGTAGATTTAGTTTGACC |
| **U-14215-13231** | TCGGTTGTATTTGGGGCGCGAGCTGAAAAGGT |
| **U-15026-13030** | CCAGCGATTTGGGAATTAGAGCCAGTTAGCAAGGCCGGAAACG |
| **U-15040-17039** | AAAAGGGCCATATAAAAGAAACGCGAATACCC |
| **U-15053-13055** | TCAACCGATTGAAGGTGAATTATCACCGATAGCAG |
| **U-15064-16064** | AGGGAGGGAAGGTAAATATAGCAAACGTAGAAAATACA |
| **U-15120-13121** | TTGCCATCAAGTGTAGCGGTCACGCCACCACACC |
| **U-15176-18176** | AGCCGGCGAACCCTAATAAATCGGATAGCCCG |
| **U-15201-13207** | ATACTTTACCAAAAACATTATGTTTAGCTATATTTTC |
| **U-15233-17232** | CAAGGATGAAAGGCCGGAGACAGGCTATTTTT |
| **U-15248-13248** | TTAGAACCAGCAAAATTAAGCAATTTCTACTAA |
| **U-16096-15103** | CTCAGAATTTTCGGTCATA |
| **U-160110-15119** | TCAGAGCGCCCCCTTATTAGCGT |
| **U-160135-18136** | GCCGCCACAGGGCGAAAAACCGTCAAGAGTCC |
| **U-160151-14152** | AACCGCCTCACCGGAACCAGAGCCAAAGCGAA |
| **U-160167-18168** | CCCGATTTAGGTGCCGTAAAGCACAGATAGGG |
| **U-160215-14216** | CCATCAATTGCGGGAGAAGCCTTTAAGCTAAA |
| **U-17040-1939** | AAAAGAACACCGAAGCCCTTTTTAGCGCTAAT |
| **U-17057-1552** | TAAGACTTACATAAAGGTGGCAAGACAT |
| **U-17064-1864** | CCTTATTACGCAGTATGTTAGAGCAAGAAACAATGAAA |
| **U-17108-16111** | CACCAGAGCTCCAACGTCAACCTCAGAACCGCCACCC |
| **U-17201-15200** | ATAAATTATGATATTCAACCGTACCCTGTA |
| **U-17233-19232** | GAGAGATAGCATGTCAATCATATATTGTAAAC |
| **U-17248-15247** | GCTATCAGAGATTCAAAAGGGTGAAAAAATTT |
| **U-18031-16026** | AGAAAAGTAAGCAGATAGCAGGAAACGCAATAATAACGAAAGAC |
| **U-18096-16097** | GGTTGAGCCGCCACCAGAACACCACC |
| **U-18135-20136** | ACTATTAAATAAATCCTCATTAAACGTTCCAG |
| **U-18151-16152** | TTTGGAACTATCATTTTGGGGTCGACCACCGG |
| **U-18167-20168** | TTGAGTGTCAAAATCCCTTATAAAGTGGTTCC |
| **U-18215-16216** | GTTGATAAAATGCCGGAGAGGGTATCAAATCA |
| **U-19040-21039** | ATCAGAGAAGAGAATAACATAAAAAAATAAAC |
| **U-19057-17056** | CAAGAATTAGCAATAGCTATCTTTGGCATGAT |
| **U-19064-20064** | TGAGTTAAGCCCAATAATATTTAACGTCAAAAATGAAA |
| **U-19113-17107** | GATATTCACAAACAAAGAACGTGGACCGCCGCCAGCATTCCAC |
| **U-19176-22176** | TCAAAAGAGTTTGATGAAAATCCTTTGCGTAT |
| **U-19201-17200** | AGATTGTTCAGAAAAGCCCCAATCTAGCTG |

| | |
|---|---|
| **U-19233-21232** | GTTAATATTCGCGTCTGGCCTTCGAACAAACG |
| **U-19248-17247** | AAATTCGCCGGTAATCGTAAAACTCTACAAAG |
| **U-19256-18256** | ATTAAATTTTGTTAAATCAGGCAAACAAGAGAATCGATGAA |
| **U-20012-21015** | GGGAGAATTAACTGCGTCTTTCCAGAGCC |
| **U-20031-19031** | ACAGGGAAGCGCATTAGTCAGAGGGTAATTGA |
| **U-20096-18097** | TGGTAAGGCAGGTCAGACGGACAGGA |
| **U-20135-22136** | TAAGCGTCCTGAGAGAGTTGCAGCGAGACGGG |
| **U-20151-18152** | GAATTTACGCCAGAATGGAAAGCGTGTTCCAG |
| **U-20167-22168** | GAAATCGGTTTGCCCCAGCAGGCGTGGGCGCC |
| **U-20215-18216** | AGCTTTCAATAAGCAAATATTTAAGTACCCCG |
| **U-21040-23039** | AGCCATATTTTGCACCCAGCTACATCCGGTA |
| **U-21057-19056** | CAATCCAATAGCAGCCTTTACAGGATAACCCA |
| **U-21064-22064** | AATAAGAAACGATTTTTTGGAGGTTTTGAAGCCTTAAA |
| **U-21124-19112** | GGCCATACATGGCTTTTGATGATACAGGATTGGCCTT |
| **U-21201-19200** | ACAACCCTCAACATTAAATGTGAAACAGGA |
| **U-21233-23232** | GCGGATTCGACAGTATCGGCCTCGCCATTCGC |
| **U-21248-19247** | TGGGATAGACGCCATCAAAAATAATTTTGTTA |
| **U-22096-20097** | CAGTTATAAGTTTTAACGGAGTGTAC |
| **U-22121-21123** | CCTTCAGTAACAGTGCCCGGTCAGTGCCTTGACCGCCT |
| **U-22135-24136** | CAACAGCTCATGAAAGTATTAAGAAGCGGGGT |
| **U-22151-20152** | TCACCAGTAAGCGGTCCACGCTGGCAGTCTCT |
| **U-22167-24168** | AGGGTGGTGGCCAACGCGCGGGGAGCTGCATT |
| **U-22215-20216** | CGCACTCCGTCGGATTCTCCGTGGCTGTAGCC |
| **U-23040-25039** | TTCTAAGAAGAACAAGCAAGCCGTTACGAGCA |
| **U-23057-21056** | GTTTTAGTCAAGATTAGTTGCTATATTTATCC |
| **U-23064-24064** | CGAACCTCCCGACTTGCGGAAGAACGGGTATTAAACCA |
| **U-23176-25183** | GAGGCGGTTCGTGCCAGAAACCTG |
| **U-23201-21200** | TGGTGCCAGCCAGCTTTCCGGCAGCGAGTA |
| **U-23233-25232** | CATTCAGCTGCAAGGCGATTAAGCATGCCTGC |
| **U-23248-21247** | ACTGTTGGCAGTTTGAGGGGACGAGACCGTAA |
| **U-24031-23031** | TTTTATTTTCATCGTAGGAATCAAATCAGATATAGAAGGCTT |
| **U-24096-22097** | TGATATATGCCCCCTGCCTGTATAAA |
| **U-24111-22122** | TGCCGTCGATTTCGGAACCTATTATTCTGAAAGATTGC |
| **U-24135-26133** | TTTGCTCAAGGTTTAGTACCGCCACGCCACCCTCA |
| **U-24151-22152** | TTAGGATTGGCTGAGACTCCTCAATTTTCTTT |
| **U-24167-26168** | AATGAATCCCCGCTTTCCAGTCGGGAGTGAGC |
| **U-24215-22216** | CGCCAGGGGGAAACCAGGCAAAGCAGGAAGAT |
| **U-25040-26031** | TGTAGAAAATGTTCAGCTAATGCAG |
| **U-25057-23056** | AATCGGCAGTACCGCACTCATCGACGCGAGGC |
| **U-25064-26065** | TGTCTTTCCTTATCATTCCAAAGTAATTCTGTCCAGA |
| **U-25116-24112** | ACCGTACTCAGGGTACCAGGCGGATAAG |
| **U-25201-23200** | AACGACGTTTTCCCAGTCACGAACCGCTTC |

| | |
|---|---|
| **U-25233-26216** | AGGTCGACCGCTCACAATTCCACACAACATA |
| **U-25245-23247** | GAGGATCCCCGGCGAAAGGGGGATGTGGCTGCGCA |
| **U-26030-25031** | AACGCGCCTGTTTATATATCCCATCCTAATT |
| **U-26064-25056** | CGACGACAATAAACAACCCAATCAAT |
| **U-26103-24097** | TATAAAGTACCGAAAGTATAGCCCGGAGAGGGT |
| **U-26132-28128** | GAACCGCCTTCGAGCATTACCCTCAGAGCTCCACAGA |
| **U-26167-27160** | TAACTCACATTAATTGAGGGATAGC |
| **U-26207-25200** | AAGCATAAAGTGTAACGTTGTAA |
| **U-26215-24216** | CGAGCCGGGCCAGTGCCAAGCTTGTTGGGTAA |
| **U-26276-25244** | ATCATGGTCATAGCTGTTTCCTGTGTGAAATTGTTATCTCTA |
| **U-27091-29097** | AACAACGCCAACAAATCGCCATATTTATAAAGC |
| **U-27113-25115** | GGCAGAGGCCAGTAATAAGAGAAAATAGGTGTATC |
| **U-27144-24152** | TCATTTTCCGTAGAACCCCTCTGCGCTCACTGGAGAAGGA |
| **U-27161-29159** | AAGCCCAGTCACCAGTACAAACTGTTAGTAA |
| **U-28127-30128** | CAGCCCTCTAACGATCTAAAGTTTAAAAGCCT |
| **U-28175-26176** | TGAGTTTCATAGGAACCCATGTACTGCCTAAT |
| **U-29098-31097** | CAACGCAATTCTTACCAGTCTGACCT |
| **U-29112-27112** | AGGTAGCGATAGTGCTTAATTGAGTGTAATTTA |
| **U-29145-27143** | TCCAGACACAACGCCTGTAGCATCACCACCC |
| **U-29160-31160** | ATGAATTTAGTTTCAGCGGACCGGTAAACAGTG |
| **U-30127-32128** | GTTTAGTAGACCGTGTGATAAATACTCCAAAA |
| **U-30175-28176** | CTTTCAACTCTGTATGGGATTTTGCGTAACAC |
| **U-31098-33097** | AAATTTTTAATTTCATCTTAAGAACG |
| **U-31112-29111** | GAAATACCTCATATGCGTTATACATCAACAGT |
| **U-31145-29144** | ATAAGAAAATCATAATTACTAGATGTCGTCTT |
| **U-31161-33160** | AGAATAGTTTTTTCACGTTGAAATAAACAGCT |
| **U-32127-34128** | GGAGCCTTGGTTTATCAGCTTGCTTATATAAC |
| **U-32175-30176** | AATAATAAAAAGGAACAACTAAAGCTAAACAA |
| **U-33098-35097** | CGAGAACAATCGCAAGACAATTAAGA |
| **U-33112-31111** | CAAGTATCTAATTATATATTTTAGAATGGTTT |
| **U-33144-31144** | GAATTTCTATCTCCAAAAAAAAGGAGGCGTTAA |
| **U-33161-35160** | TGATACCCCCACGCATAAAACCTTTTTTCCGA |
| **U-34127-36128** | TATATGTAATTTATCAAAATCATAGACAGCAT |
| **U-34175-32176** | AACCATCGGATAGTTGCGCCGACAGAATTGCG |
| **U-35098-37097** | CGCTGAAGCGATAGCTTAGATTAATT |
| **U-35112-33111** | AATAGTGAAATGCTGATGCAAATCAACTTTTT |
| **U-35144-33143** | AGACTACCCCGGCTTAGGTTGGGTTTCGAGGT |
| **U-35161-37160** | TATATTCCTTTTGCGGGATCGTCTCATGAGGA |
| **U-36127-38120** | CGGAACGAGGCTACAGAGGCTTTGATCAATATATGTGAGT |
| **U-36175-34176** | AAAGGCCGGGTCGCTGAGGCTTGCATGACAAC |
| **U-37098-39097** | AATTTTGTAAATCGTCGCTGATGATG |
| **U-37104-35111** | CCCTTAGAATCGCAACGGGTACTTGAAAACATGAAGAGTC |

| | |
|---|---|
| **U-37145-35143** | GACTTTTACCCTCAGCAGCGAAAGGTCTGAG |
| **U-37161-39162** | AGTTTCCAAGGCACCAACTTTAACCTTTCTAAAA |
| **U-38119-39120** | GAATAACCTTGCTTCTACATCAAGAAAACAAAA |
| **U-38175-36176** | CCACTACGATTAAACGGGTAAAATAGGGAGTT |
| **U-39121-41109** | TTAATTACGGAGATTTGTTCATTTCAATTACAGAGGC |
| **U-39147-37144** | AATTATGGAAACAGTACATAAAGGACTAAA |
| **U-39163-40150** | CGAAATGACCCCCAGCGATTATA |
| **U-40135-39146** | AAGTACAACATTTAACAATTTCATTTG |
| **U-40149-42154** | CCAAGCTCCGCGACCTGCTCCATGTTACTTGGGAACCGTACAGT |
| **U-41110-42111** | GAATTATATCATCGCCTGCAATAACGGATTCGCCT |
| **U-42110-44120** | GATTGCTCGTAAAACAGAAATAAAGAAATTGCTTGACAA |
| **U-42135-40136** | GGGAGAAAATAAATTGTGTCGAAAGCGAAACA |
| **U-42153-44156** | AACAGTACCTTAACGTCAGATGAATAAACTGACCGACCAGGCGCAT |
| **U-44119-45120** | GAAATGGAAGGGTTAGATACTTCTGAATACCGG |
| **U-44135-42136** | AGAGTAATCGTAGATTTTCAGGTTTTTACATC |
| **U-44155-46163** | AGGCTGGCTGACAACAAAGCTGCTCATTCAGTGAATGTAGT |
| **U-45121-47108** | ATATTCATGATTATCAGATGATGGCAATTCAAACGT |
| **U-46135-44136** | CATATTCCTTACCCAAATCAACGTCTTCATCA |
| **U-46162-48152** | AAACACCAGAAGGAGCGGATGCGGAACAAAGAAACTATGCGAT |
| **U-47109-48124** | TATTAATTTTAAAAGTTTGGTCA |
| **U-48123-49120** | GGACGTTTACAAACAATTCGAAGTATTAGACTTGGGA |
| **U-48135-46136** | TTATACCAAGTAACATTATCATTTATTATCAT |
| **U-48151-50152** | TTTAAGAAGAACTAACGGAACAACTTCAACTA |
| **U-49121-50110** | AGAAAAAAACAACTAATAGATTAGA |
| **U-50109-0120** | GCCGTCTTGAAAGGAATTGAGGAAGGTTATCTCAAATA |
| **U-50135-48136** | GGAGCACTTCTACGTTAATAAAACCTGGCTCA |
| **U-50151-0144** | ATGCAGATAAAAGGAATTACGAGGAATGAAAAATCTAAAG |
| **U-50162-47167** | ACAATTATTACAATTACCTTTGGGCTT |
| **U-1177-51170** | AGCGAGAACCCTCGTTTACCAGAGAG |
| **U-14262-15276** | AAGAATTCTCATATATTTTAAATGCAAT |
| **U-21256-20256** | GTCACGTTGGTGTAGATGGGCCTCATTTTTAACCAATAGGA |
| **U-24276-25265** | TCTTCGCTATTACGCCAGCTGGGTACCGAGC |
| **U-40174-38176** | TCATCTTGAGGCAAAAGAATACAACGTAATG |
| **U-42175-41183** | AATCATAAAGCCGGAACGAGGCGC |
| **U-44175-42176** | GGTGTACAAACTTTGAAAGAGGACAGACGGTC |
| **U-46175-44176** | CAGAACGAAAGGCTTGCCCTGACGAGATGAAC |
| **U-50183-48176** | CAGTTGAGATTTAGGAAGGTAGAAAGATTCATAACTTTAA |
| **U-13000-15012** | AGCACCATTACCACAAAATCACCAGTAATCAATAGAAAA |
| **U-15013-16007** | TTCATATGGTTTAACCACGGAATAAGTTTATT |
| **U-16006-18000** | TTGTCACCAGAAGGAAACCGCGAACAAAGTTAC |
| **U-21016-22011** | TAATTTGCCAGTTACAAATTTTATCCTGAATCTTACC |
| **U-22010-25005** | AACGCTAACGACCAATAGCAAGCATTACCGCGCCTGAAC |

| | |
|---|---|
| **U-25006-26000** | AAGAAAAATACAACAATAGATAAGTC |
| **U-39098-41097** | AAACAACTGAGCAAAAGAATACAAAA |
| **U-41098-43097** | TCGCGCTTGAATACCAAGTAAAATTA |
| **U-43098-45098** | TTTGCAAACCTACCATATCGATTGTTT |
| **U-47091-49097** | AATCCTTTGCCCGACAACTCGTATTATTTGAGG |
| **U-49098-51097** | ATTTAGAATAGATAATACAGGCAAAT |
| **U-51098-1097** | CAACAGTATCTGGTCAGTTGCAGAAG |
| **U-17256-16256** | GTCATTGCCTGAGAGTCTGGAGCCTGAGTAATGTGTAGGTAA |
| **U-23256-22256** | GAAGGGCGATCGGTGCGGGCCGCATCGTAACCGTGCATCTGC |
| **U-48175-46176** | TCATTGTGGAGATGGTTTAATTTCAGAAACAC |
| **1L-25266-40175** | TCGAATTCGTACTAAAACAC |
| **1L-51171-14263** | CAACACTATCATAATACAGGCAAGGCA |
| **CB-45099-20013** | GGATTTCAATATAATCCTGAACACCCTGAACAAAGAC |
